# The structure of hippocampal CA1 interactions optimizes spatial coding across experience

**DOI:** 10.1101/2021.09.28.460602

**Authors:** Michele Nardin, Jozsef Csicsvari, Gašper Tkačik, Cristina Savin

## Abstract

Although much is known about how single neurons in the hippocampus represent an animal’s position, how cell-cell interactions contribute to spatial coding remains poorly understood. Using a novel statistical estimator and theoretical modeling, both developed in the framework of maximum entropy models, we reveal highly structured cell-to-cell interactions whose statistics depend on familiar vs. novel environment. In both conditions the circuit interactions optimize the encoding of spatial information, but for regimes that differ in the signal-to-noise ratio of their spatial inputs. Moreover, the topology of the interactions facilitates linear decodability, making the information easy to read out by downstream circuits. These findings suggest that the efficient coding hypothesis is not applicable only to individual neuron properties in the sensory periphery, but also to neural interactions in the central brain.

---

The dual role of the hippocampal formation in memory (1, 2) and spatial navigation (3, 4) is reflected in two distinct views on hippocampal coding: the place field view (5, 6) that reduces the encoding of spatial information to tuning properties of individual neurons, and the ensemble view (7, 8) that focuses on subsets of units that are co-activated together as the substrate for memory (9). Recent results blur the line between the single cell and the population perspective (10), revealing that properties of individual neurons only partially explain the circuit’s contribution to spatial encoding. Interactions between neurons shape collective hippocampal activity (11) and contribute to the spatial representation. Disrupting correlations between neurons leads to decreased decoding accuracy, in particular in CA1 (10). It remains unclear how experience shapes the organization of cell-to-cell interactions and what effects such changes may have on the encoding of spatial information in CA1 populations.

Experience affects the properties of single cells in many ways. While reliable position-dependent spiking is detectable after a few minutes during the very first exposure to a novel environment (12, 13), the responses to a familiar environment show several systematic differences, including a reduction in overall firing, sharpening of tuning functions and sparsification of responses (14). In parallel, inhibition is weak in novel environments, transiently opening the gate for circuit reorganization via plasticity (15), but it subsequently increases with experience (15–17). From the perspective of the local circuit, the main afferents to CA1 (MEC and CA3) are initially noisier (18, 19) and have weaker spatial tuning, which improves with familiarity (13, 20, 21). Since CA1 needs both inputs for detailed spatial representation (22, 23), these results suggest that the CA1 circuit is potentially in a different dynamic regime in novel versus familiar environments, with distinct local circuit interactions and population coding properties.

Correlations among pairs of hippocampal neurons arise as a result of two effects: their spatial tuning overlap (i.e. *signal* correlations), and internal circuit dynamics (i.e. *noise* correlations). Since they reflect local circuit interactions, noise correlations should depend on changes in input statistics, and be reorganized by experience. From a neural coding perspective, the structure of neural correlations can radically affect the amount of information that a population carries about stimuli (here, the animal’s position) and the complexity of the readout (24, 25). While noise correlations are generally considered to be an obstacle to optimal information coding and transfer, especially in sensory areas (26, 27), there are scenarios where they can improve the quality of the overall population output (28–31), which might be relevant for the hippocampus.

Unlike sensory areas, where stimulus repeats make the estimation of noise correlations relatively straightforward, measuring circuit interactions and their contribution to spatial coding in the hippocampus is fraught with technical difficulties. In a two dimensional environment, the lack of stimulus repeats renders traditional approaches for estimating noise correlations inapplicable. Moreover, well documented circuit level oscillations (32, 33) act as global sources of co-modulation that obscure the fine structure of pairwise neural co-variability. The key challenge is to partition total neural covariability into an explainable component, driven by position and oscillations, and unexplained, or ‘excess’ correlations, which capture local interactions.

Here we take advantage of the maximum entropy framework to develop a new statistical test for detecting excess correlations without stimulus repeats, and explore their significance for the encoding of spatial information in CA1. Our method allows us to robustly detect network interactions by comparing hippocampal responses against a maximum entropy null model (34) that optimally captures the cells’ place preference and population synchrony (35). When applied to CA1 tetrode recordings from rats during open field exploration in familiar and novel environments, our analysis detected structured excess correlations preferentially between principal cells with similar place selectivity and arranged into networks with high clustering coefficients. These highly structured excess correlations optimize the encoding of spatial information and facilitate its downstream readout in both the familiar and novel environment, with differences reflecting the different signal-to-noise ratio of spatial inputs in both environments. Taken together, our results suggest that CA1 local circuitry readjusts to changes in its inputs so as to improve population-level stimulus representation, in line with efficient coding predictions (29).

## Results

### Detecting interacting cells

To investigate functional connectivity between CA1 neurons and its role in spatial information coding, we devised a procedure to infer cell-cell interactions from simultaneous tetrode recordings of hundreds of isolated units in dorsal hippocampus of behaving rats.

Our approach starts by constructing a null model for population responses that exactly accounts for the measured spatial selectivity of each recorded neuron as well as for the moment-to-moment measured global neural syncrhony, but is otherwise as unstructured as possible (Fig. 1A). This null model is formally a maximum entropy model (see Methods) from which surrogate neural rasters can be sampled (34). For every cell pair, the model predicts the expected distribution of pairwise correlations against which the measured total correlation for that pair can be tested for significance; we report as “excess correlation” *w* the (normalized) amount of total correlation that is not not explained by the null model. We declare cell pairs with significant excess correlation to be “interacting,” likely due to specific recurrent neural circuitry. Because our approach explicitly discounts for correlations arising from overlapping place fields and global modulation (e.g, due to locking to the underlying brain oscillations or influence of behavioral covariates such as running velocity), it differs from previous attempts to use correlations to probe the intrinsic network mechanisms (36).

**Figure 1.**
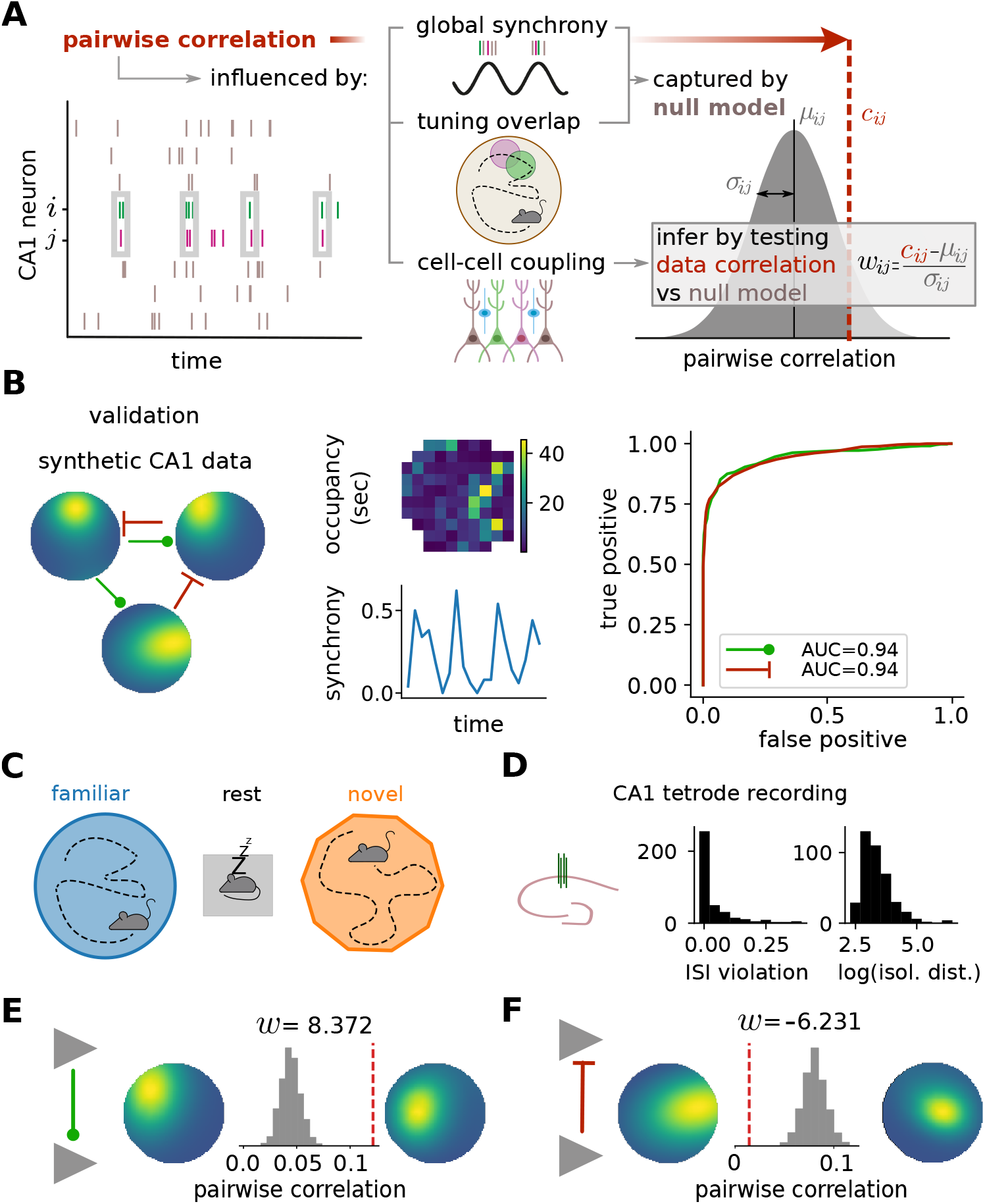
Detecting network interactions among hippocampal CA1 cells. **(A)** Method schematic. A null model for population responses takes into account the inferred place field tuning of each cell and the moment-to-moment global synchrony, but is otherwise maximally unstructured. For each cell pair, this model predicts a null distribution for (total) pairwise correlation (gray distribution), which is compared to the correlation estimate from data (dashed red line). The normalized discrepancy between the data correlation *c*_*ij*_ and the null model expectation *µ*_*ij*_ for a pair of neurons (*i, j*) is referred to as “excess correlation”, *w*_*ij*_, and serves as a proxy for direct cell-cell interaction. **(B)** Method validation on synthetic data. Detection accuracy is assessed using simulated data with known interactions (left), which matches real data with respect to spatial occupancy (top, middle) and observed synchrony indices (bottom, middle), for an example 20-minute exploration session. Receiver-operator characteristic (ROC) shows the probability of correctly detecting positive (green) and negative (red) interactions for different detection thresholds (right). **(C)** Experimental paradigm. Animals explore a familiar environment, then rest in a sleep box, after which they explore a novel environment (20–40 minutes for each condition). **(D)** Neural recordings. Left: neural activity was recorded using tetrodes implanted in the dorsal CA1 area of the hippocampus. Middle: distribution of ISI violation scores after spike sorting for the data included in the analysis. Right: same for the Isolation Distance measure. **(E,F)** Example pair of pyramidal cells with significant positive (E) and negative (F) excess correlation *w* (gray histogram – distribution of correlation coefficients derived from the null model; red dashed line – measured raw pairwise correlation).

We validated our detection method by constructing a synthetic dataset of spiking CA1 neurons whose responses were modulated by the position of an artificial agent and by an assumed network of interactions (see Methods). We ensured that the synthetic data matched the synchrony and the highly irregular occupancy observed in a real 20-minute exploration session. Interactions identified by our method strongly overlap with the ground truth, as measured by the area under the receiver operating characteristic (Fig. 1B). The inferred excess correlations were well aligned with the ground truth (Fig. S1A). We did not find any tendency of cells that are more (or less) similarly tuned to show higher (or lower) inferred *w*_*ij*_ s (Fig. S1B).

We next analyzed CA1 tetrode recordings of six rats exploring familiar and novel 2D environments separated by a short period of rest (Fig. 1C) (37, 38). Putative units were filtered by using several clustering quality measures (39–41) to ensure that they were well isolated (Fig. 1D, see Methods). To avoid confounds due to changes in firing rate, we retained only cells active in both environments (> 0.25 spike/sec) (14). Considering only pairs of cells recorded on different tetrodes, our final dataset includes a total of 9511 excitatory-excitatory (EE), 7848 excitatory-inhibitory (EI), and 1612 inhibitory-inhibitory (II) pairs. We detected both positive and negative excess correlations among cell pairs (Fig. 1E,F). Interestingly, cell pairs with negative excess correlation can have positive total correlation (Fig. 1F), corroborating the idea that the network circuitry can strongly affect coordinated spiking activity in the hippocampus.

### Interaction networks in familiar and novel environments

What is the structure of the inferred interaction network? We set the threshold to declare a cell pair as interacting at |*w*| > 4.5 (corresponding to a p-value cut of *p* = 0.05 prior to Bonferroni correction for multiple comparisons; see Methods). We first report a generally sparse interaction network in the excitatory-excitatory (EE) subnetwork, with ∼ 5% of analyzed pairs showing significant interaction; this coincidentally implies that our null model accounts for most of the observed correlation structure. The fraction of interactions is larger among excitatory-inhibitory (EI) cell pairs, where, as expected, negative interactions dominate; the fraction is highest at ∼ 30% among positive interactions in the inhibitory-inhibitory (II) subnetwork (Fig. 2A).

**Figure 2.**
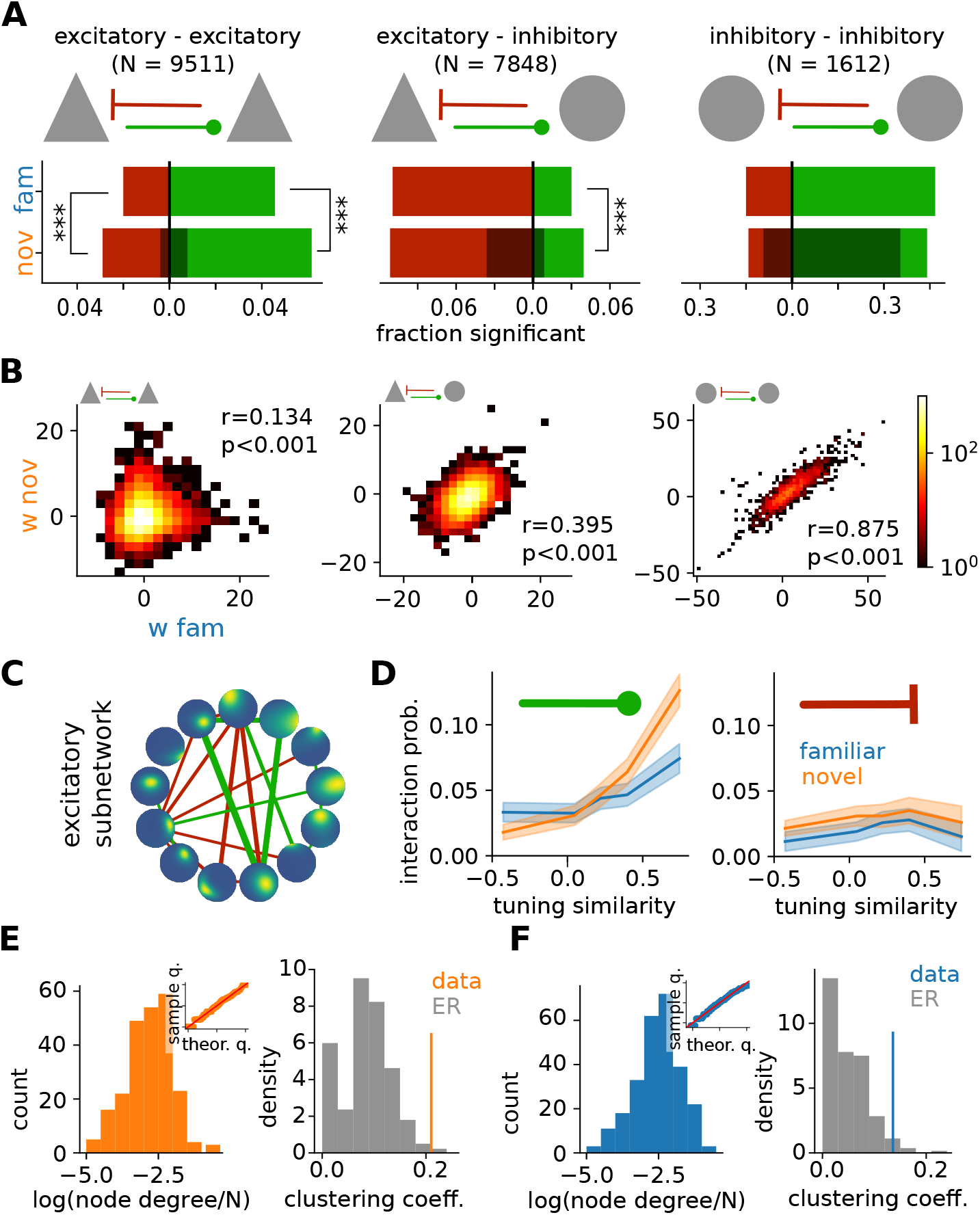
Network interactions in familiar and novel environments. **(A)** Summary of cell-cell interaction results for different cell types (triangle – pyramidal cell, circle – putative interneuron), positive (green) and negative (red) excess correlations, for both the familiar (top row, blue) and the novel (bottom row, orange) environment (stars – significant difference under binomial test at *p <* 0.001). Shaded regions mark the fraction of interactions detected in the familiar environment that remain in the novel environment. **(B)** Paired comparison (colormap – pair density) between excess correlations *w*_*ij*_ detected in familiar vs. novel environment. **(C)** Example of an estimated excitatory subnetwork. Circles show the place field selectivity of each neuron, with edges showing significant cell-cell interactions (green – positive; red – negative excess correlations); line thickness corresponds to interaction strength. **(D)** Left: interaction probability in the excitatory subnetwork increases with place field overlap (“tuning similarity”) for positive interactions (blue – familiar environment; orange – novel environment; shaded area – 99th percentile confidence interval for the mean). Right: analogous plot for negative interactions. **(E)** Left: distribution of log node-degree of E cells normalized by the total number of E cells in each session, for the novel environment. Inset: quantile-quantile plot comparing this distribution to the normal expectation. Right: excitatory subnetwork has a significantly higher clustering coefficient (orange line – data) compared to the expected distribution for an Erdos-Renyi (ER) network with a matched connection density. **(F)** Same as (E), but for the familiar environment.

We next focused on interaction changes induced by a switch from familiar to novel environment (Fig. 2A). We observed a significant increase in EE interactions, possibly due to decreased inhibition during novelty (15, 17), which enhances learning and promotes plasticity (42–44). We indeed found putative inhibitory cells to be less synchronous and slightly less active in novel environments (Fig. S2B,D), in line with previous findings (16), while excitatory neurons were more synchronous but did not differ in terms of their average firing rates (Fig. S2A,C). Circuit modifications during spatial learning are known to originate in altered spike transmission among connected excitatory and inhibitory neurons (45, 46). Consistent with this, we found an increase in positive EI interactions, while their negative counterpart remained unchanged. This increase could not be attributed to increased reliability of monosynaptic EI connections (Fig. S3), especially since cell pairs on the same tetrode were excluded (47). We did not observe significant changes in the number of II interactions.

How conserved are individual network interactions across consecutive environments? The largest overlap in detected interactions was found in the II subnetwork, where 77.5% of interactions were preserved, preferentially among cell pairs with similar theta sensitivity (Fig. S4D; (48)). EI interactions, especially inhibitory, also showed substantial overlap (31.1%); the correlation with theta selectivity was small but significant (Fig. S4D). The overlap was weakest (16.8%) in the EE subnetwork; no correlation with theta selectivity was observed (Fig. S4D).

All reported overlaps were statistically significant under a permutation test (1000 random shuffles of cell labels; *p <* 10^−3^ for all subnetworks). Significance was confirmed by comparing the Jaccard similarity of the adjacency matrices of familiar and novel subnetworks against the null distributions constructed from Erdos-Renyi graphs with matched numbers of vertices and edges (1000 ER graphs; *p <* 10^−3^ for II and EI subnetworks, *p* = 0.009 for EE).

The similarity of interaction networks across the two environments extends beyond the binary presence / absence of significant interactions. Figure 2B compares the strength of excess correlations, *w*, in familiar vs novel environment for EE, EI, and II cell pairs. For all subnetworks, *w* are significantly correlated across the two environments, with the reported correlation strength related to the network overlap (Fig. 2A). Taken together, these findings corroborate the idea that hippocampal remapping across environments is not random (49), also at the level of cell-cell interactions.

Because spatial information is encoded predominantly by pyramidal cells (50, 51), we analyzed the EE subnetwork in detail (Fig. 2C). Our key statistical observation is shown in Fig. 2D: interaction probability increases nonlinearly with place field overlap for positive interactions, and is roughly constant for negative interactions. In the novel environment, the excitatory interaction probability increases ∼ 3-fold over the observed range of place field overlap. In the familiar environment, the modulation with place field overlap is less pronounced, possibly indicating a shift towards a more decorrelated representation of space (14).

We further characterized the topology of familiar and novel excitatory networks. The node degree appears to be log-normally distributed in both environments, with clustering coefficients that are significantly higher than expected from matched Erdos-Renyi graphs (Fig. 2E,F). This effect was more pronounced during novelty (Fig. S5A), in line with recent reports (36). Accordingly, interacting excitatory triplets were over-represented, more strongly so in the novel environment (Fig. S5C). Finally, we found a linear relationship between the log-number of nodes and the shortest path length (Fig. S5B), which is a strong fingerprint of small-world notworks (52).

### Effects of network interactions on spatial coding

To explore how the network structure affects spatial information encoding at the population level, we constructed a statistical model of interacting excitatory cells responding to spatial inputs (Fig. 3A). Our model, a version of pairwise-coupled, stimulus-driven maximum entropy distribution over binary spiking units (see Methods, (53)) allows us to vary cell-cell excess correlations (to study the effect of network topology and interaction strength) as well as the strength of the spatial inputs (to study the effect of novel vs familiar environment), while maintaining a fixed average firing rate in the population. For tractability, we simulated populations of 50 place cells. Our model is thus not an exact fit to data or at-scale model of the real hippocampal population; rather, we are looking for qualitative yet clear signatures of spatial coding at the population level that could be compared between the data and the model.

**Figure 3.**
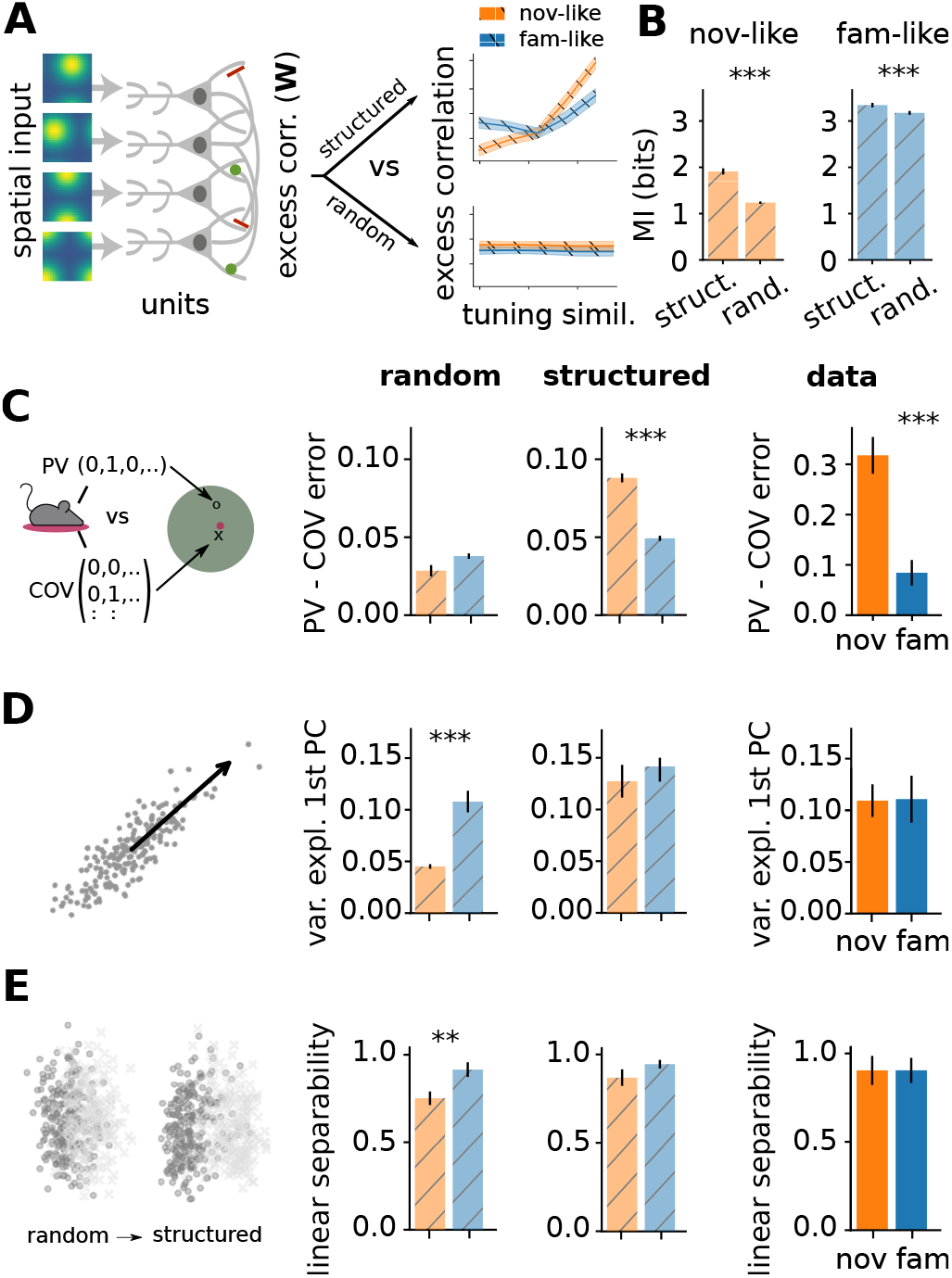
Effects of network interactions on spatial encoding. **(A)** A schematic of the circuit model with variable excess correlations (see Methods). Two connectivities are compared: “structured” (mimicking the inferred excess correlation vs tuning similarity relationship) vs. “random”. **(B)** Estimated spatial information (MI; error bar – 99-th percentile CI for the mean) using structured and random interactions, in the novel-like and familiar-like scenario (see text). Structured interactions significantly increase the spatial information (*p <* 0.001 (***) or *p <* 0.01 (**) under a non-parametric Mann–Whitney U-test). **(C)** Improvement in decoding performance by taking into account co-variability of cells (“COV” decoder) relative to a simple population vector (“PV”) decoder, evaluated on 4 · 10^4^ samples). The improvement is significantly higher in the novel environment on structured network and on real data, but not on the random network (error bars and significance tests as in B). **(D)** Fraction of variance explained by the first principal component of population vectors for 10^3^ random pairs of locations in the maze. The fraction is unchanged between the novel and familiar environments on structured network and on real data, but differs significantly on the random network (error bars and significance tests as in B). **(E)** Linear separability measured as SVM classification accuracy of random pairs of stimuli (trained on 1000 pairs of same vs. different positions). The separability is unchanged between the novel and familiar environments on structured network and on real data, but differs significantly on the random network.

Using this setup, we contrasted spatial coding in two networks which were identical in every respect except for their excess correlations pattern. Interactions in the “structured” network followed the relationship between place field overlap and excess correlation *w* observed in real data; interactions in the “random” network were drawn from the same data-derived distribution for *w*, but did not follow the relationship with place field overlap (Fig. 3A). For each of the two choices, we further simulated the effects of familiar vs. novel environment by adjusting the strength of the feed-forward spatial input: in our model, higher input strength corresponds to higher signal-to-noise ratio for the spatial drive, which is why we refer to this parameter as “input quality”. We adjusted the input quality to best resemble various marginal statistics (spatial information, place field sparsity, peak-over-mean firing values; see Methods and Fig. S6) in familiar and novel environments measured on data.

We quantified the coding performance of our networks by estimating the mutual information between population activity and location and by estimating the average decoding error. As expected, higher input quality in the familiar environment leads to overall higher information values (Fig. 3B) and lower decoder error (Fig. S7B). Less trivial are the effects of network connectivity: in both environments, structured (data-like) interactions significantly outperform random ones, with larger improvements seen in the novel environment. This suggests that network interactions among hippocampal cells adjust to maintain a high-fidelity spatial representation even when they receive lower quality, noisy inputs.

Do the structured interactions better predict other population-level aspects of the real hippocampal code relative to random ones? First, we assessed the importance of pairwise (co-firing) statistics for the decoding performance, highlighted by previous work (10). For the random network, the decoding performance improvement with co-firing statistics relative to population-vector decoding is small and comparable in novel vs familiar environment. In contrast, for the structured network and data, the improvement is significantly larger in the novel environment (Fig. 3C); the improvement reaches three-fold in novel relative to the familiar environment on real data, perhaps due to the larger population size.

Second, we assessed the effective dimensionality of the population responses to random pairs of stimuli, by measuring the fraction of variance explained by the first principal component of the relevant activity patterns (Fig. 3D). For the random network in the novel environment, this fraction is two-fold lower than in the familiar environment. In contrast, for the structured network and data, the fraction is about 0.1 regardless of the environment. Stronger and structured interactions appear to organize neural responses in the novel environment so that the code maintains a collective correlated response even when the input drive is weak.

Third, we assessed the linear separability of spatial positions based on neural population responses, a task putatively carried out by downstream brain areas. For the random network, the performance of a linear classifier trained to discriminate random positions is significantly worse in the novel environment. In contrast, the performance is restored to a high value (∼ 0.9) irrespective of the environment by data-like interactions in the structured model, matching observations on real data (see Fig. S8 for separability of positions as a function of their mutual distance).

Taken together, our results suggest an important coding role for the interaction patterns inferred in Fig. 2D and the corresponding “structured” networks explored in Fig. 3. In comparison to the random network, the data-like, structured network (i) encodes more information about position even when the input is of low quality; (ii) this information can be retrieved by utilizing co-firing statistics of multiple cells; (iii) selected collective statistics of place cell activity remain constant under change of environment. Consistent conclusions hold for the comparison between the data-like, structured network and an uncoupled population (Fig. S7).

### CA1 interactions match predictions of an optimal coding model

While Figure 3 suggests that interactions between cells self-organize to improve spatial information coding relative to a random or an unconnected (Fig. S7) network, it is not clear whether the observed organization is in any sense optimal. To address this question, we numerically optimized cell-cell interactions among a population of place cells, so as to maximize the mutual information between the population activity and spatial position (Fig. 4A). In essence, this amounts to finding “efficient coding” solutions for network structure given inputs to individual cells that are correlated due to place field overlaps (29). As before, an important control parameter is the overall magnitude (quality) of the input drive, *h*, which we now vary parametrically. Resource constraints were simulated by constraining the optimization to keep the global firing rate constant and the possible couplings bounded in |*W*_*ij*_ |*≤ w*_max_ = 1 (see Methods).

**Figure 4.**
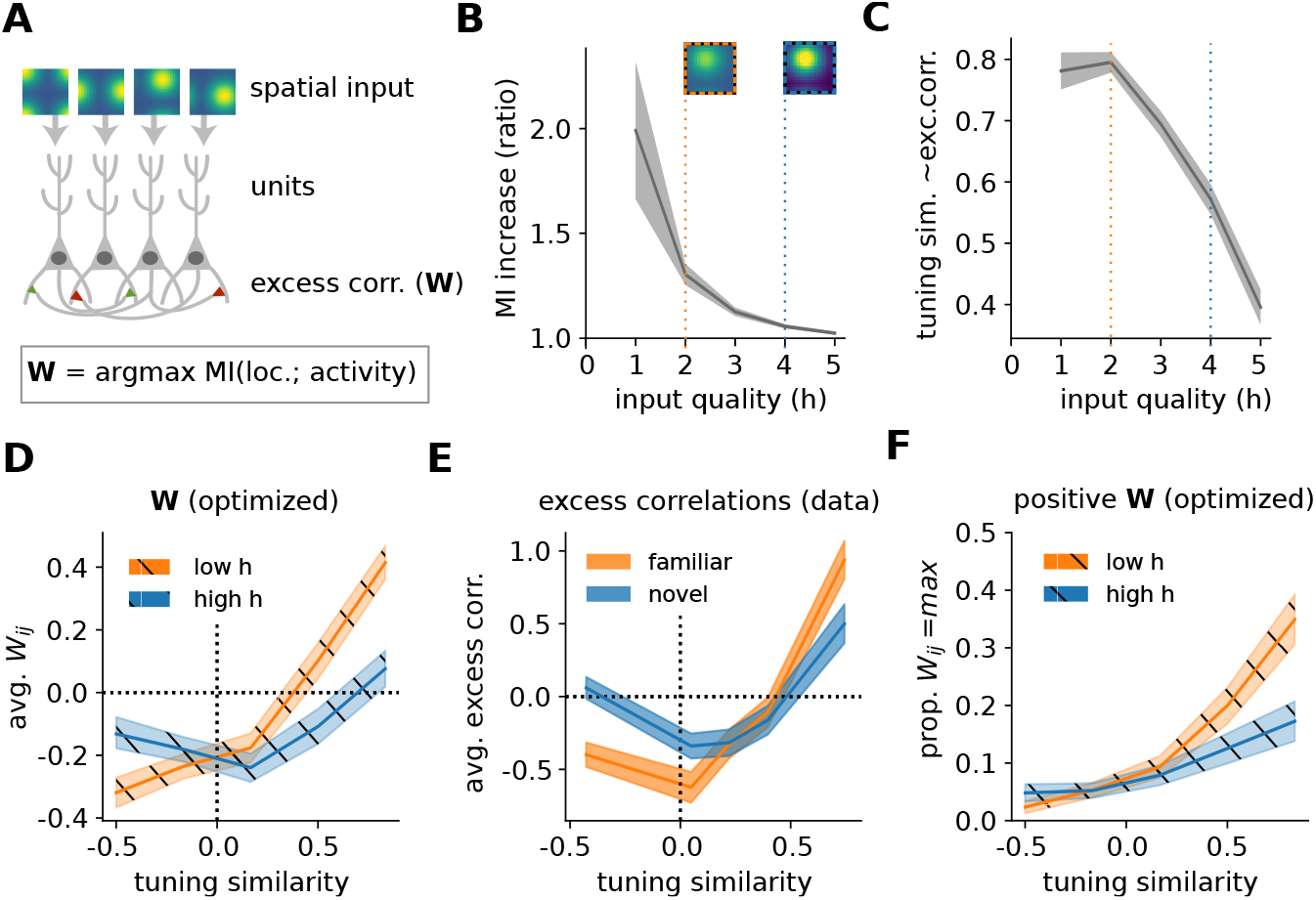
Predicted optimal network interactions. **(A)** A schematic of the circuit model. Individual neurons, which receive spatially tuned inputs (with overall strength controlled by parameter *h*), are pairwise connected with interactions **W**; interactions are numerically optimized to maximize the mutual information between spatial position and population responses while constraining population mean firing rates and |*W*_*ij*_ |*≤ w*_max_ (here, *w*_max_ = 1). **(B)** Average ratio between mutual information (MI) in optimized vs non-interacting (**W** = 0) networks. Dashed vertical lines denote two chosen input quality levels, together with firing rate map of an example cell (“low quality” *h* = 2, orange, resembling novel environment; “high quality” *h* = 4, blue, resembling familiar environment). In all simulation plots we show averages over 1000 replicate optimizations with random initial assignments of place fields (see Methods); shaded area – 95th percentile CI for the mean. **(C)** Average alignment (Spearman’s correlation) between pairwise input similarity and optimal *W*_*ij*_ as a function of input quality. **(D)** Average magnitude of optimal *W*_*ij*_ as a function of tuning similarity for the two environments. **(E)** Same as E, computed using the excitatory-excitatory excess correlations *w*_*ij*_ estimated from data. Note the vertical scale difference between (D) and (E): excess correlations *w*_*ij*_ are a statistical proxy for the true interactions *W* ; the two are expected to be correlated but not identical (cf. Fig. S1A). **(F)** Proportion of optimal *W*_*ij*_ = *w*_max_ = 1 as a function of tuning similarity.

As the input quality increases, the information gain due to optimal interactions decreases, indicating that optimization benefits novel environments (with noisy spatial inputs) more than familiar environments (with reliable spatial inputs) (Fig. 4B). We further find that an overlap in tuning similarity between two cells correlates with optimal pairwise interaction between them when input quality is low, but this correlation grows weaker with increasing input quality (Fig. 4C), consistent with theoretical expectation (29).

Does optimization predict a clear relationship between the tuning similarity and interaction strength for pairs of cells? Figure 4D shows two such relationships, for high and low input quality, predicted *ab initio* by maximizing spatial information. The optimal relationships closely resemble two analogous curves, for the familiar and novel environment, inferred from data (Fig. 4E). A similar resemblance is not observed if one maximizes spatial information carried by individual cells (Fig. S9), highlighting the importance of information coding at the population, not individual-cell, level.

As an alternative comparison to experiment we also studied the proportion of optimized couplings that reached maximal allowable strength (Fig. 4F; Fig. S10). In the data, cells are declared as interacting when their excess correlation exceeds a threshold, and so Fig. 2D represents a direct counterpart to our theoretical prediction. We observe a clear qualitative match that includes the decrease in proportion of strong couplings for familiar environments (Fig. S10). We further observe that the proportion of optimal couplings reaching the constraint *w*_max_ scales nonlinearly with the tuning similarity, as in the data; the shape of the nonlinearity depends on the imposed *w*_max_ (Fig. S11).

Even though our simulations use a coarse-grained and down-scaled model of a real neural population (precluding exact comparisons), we observe an excellent qualitative match between theoretical predictions and the data. Taken together, this opens up an intriguing possibility that network interactions in the hippocampus dynamically adapt to new environments so as to maximize the fidelity of population-level spatial representation.

### Central role for the nonlinear dependence of connectivity on tuning

So far, our analysis of data as well as of optimized networks has identified a consistent pattern: the nonlinear dependence of interaction probability on tuning similarity (Fig. 2D; 4F). Figure 3 further showed that the pattern is necessary, since coding benefits were absent in randomized networks. The key remaining question is whether the observed connectivity pattern is not only necessary, but also sufficient to convey spatial coding benefits and generate networks of a particular topology.

To address this question, we generated model networks of 50 place cells, as before, but limited their connection strengths to three possible values, {− *J*, 0, +*J*}, where *J* ∈ [0, 1] could be varied parametrically. We now used the interaction pattern of Fig. 2D as an actual *connectivity rule*: we selected 6% of pairs (as in data) to have a positive connection +*J* and connected them according to their tuning similarity as in data (Fig. 5A, “data-like”). To assess the role of the nonlinearity, we compared this with networks where the connection probability was linear in tuning similarity (“linear”) or where it was constant (“random”). In each of the three cases, a randomly chosen 3% of the place cell pairs (as in data) were connected with a negative strength, *−J*. As before, we fixed the average firing rate, and considered two levels of input quality, mimicking the familiar and novel environments (see Methods). This setup removed all structure (specifically, by making all connections have the same magnitude) except for that generated by the connectivity rule, allowing us to test for sufficiency.

**Figure 5.**
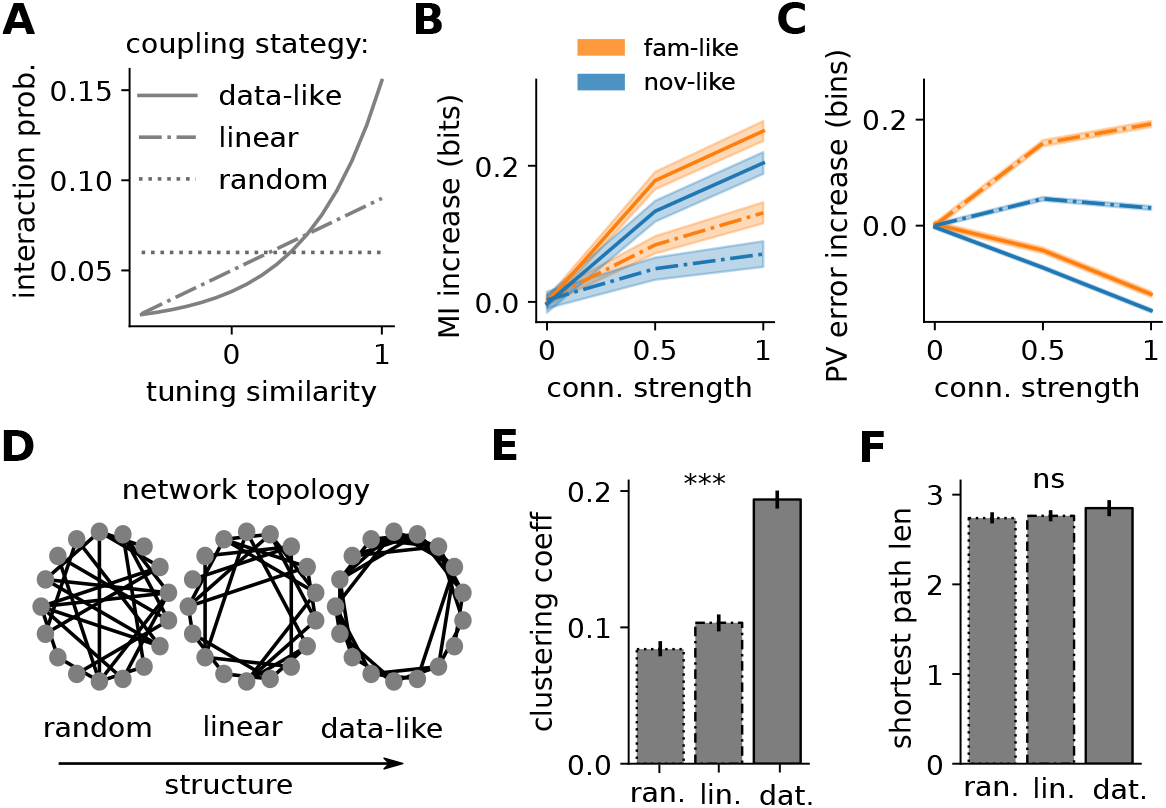
Data-like interaction pattern is sufficient to generate small-world networks with improved spatial coding properties. **(A)** Connectivity rules for positive connections in a simulated place cell network with 50 units. **(B)** Mutual information (MI) increase for data-like (solid) and linear (dashed) connectivity rule relative to the random connectivity, for familiar-like (blue) and novel-like (orange) quality input. Shaded areas show the 95th percentile confidence interval for the mean. **(C)** Average decoding error increase for data-like (solid) and linear (dashed) connectivity rule relative to random connectivity. **(D)** Example network topologies obtained by using different connectivity rules from (A). Nearby nodes have high tuning similarity. **(E)** Average clustering coefficient for the three connectivity rules from A (error bars – standard error; significance – 1-way ANOVA test, *p <* 0.001 for ***, or n.s. for *p >* 0.05). **(F)** Average shortest path length for the three connectivity rules from A (statistics as in E).

First, we find that the data-like connectivity rule consistently improves mutual information between the population responses and position for increasing *J*, especially for novel-like input quality (Fig. 5B). This improvement is larger for the nonlinear, data-like connectivity than for the linear one. Figure S13 further suggests that connectivity alone accounts for a large fraction of mutual information gain, without the need for the fine-tuning of the interaction strengths. The data-like connectivity rule also improves the performance of a simple population vector decoder relative to random connectivity, in stark contrast to the linear dependence, which performs worse than the random one (Fig. 5E).

Finally, we asked whether different connectivity rules leave a strong signature on the network topology (Fig. 5D). To this end, we randomly generated 1000 networks according to the three different rules (Fig. 5A). The average clustering coefficient was substantially higher in networks created using the data-like rule (Fig. 5E) compared to both the random and linear connectivity rules, without significantly affecting the distribution of incident edges (Fig. S12A) or the average shortest path length (Fig. 5F). Additional analysis on the clique-complexes of the connectivity graphs revealed that the 1D Betti numbers are significantly smaller for the synthetic networks generated using the data-like rule, and comparable with the data-derived networks (Fig. S12C). These analyses are consistent with the overexpression of triangles (Fig. S5) and high clustering coefficients (Fig. 2E) observed in the data-derived network. Taken together, the nonlinear, data-like connectivity rule appears sufficient to generate small-world topologies matching data across a broad panel of network metrics.

## Discussion

Statistical challenges limit our understanding of how experience shapes interactions and, consequently, information coding in a local neural circuit during animal-driven behavior. While the idea of analyzing pairwise correlations as a window into network interactions is not new (54–56), the statistical problem of separating local network interactions from other factors that drive neural correlations has remained unsolved. Previous approaches based on stimulus-averaged correlations (57), shuffles (58) or GLM model fits (59) each suffer from statistical limitations (in terms of sample efficiency, strong stationarity or other model assumptions) which limit their general applicability. For this reason, most analyses of hippocampal collective behavior rely on total correlations (36, 60). Unfortunately, these conflate changes in coding and changes in behavior: even if the representation does not change at all, a change in the animal’s behavior (e.g. with experience) would be sufficient to change collective interactions defined based on total correlations. Furthermore, well documented theta oscillations, which arise from an interplay between medial septum inputs and hippocampal subcircuits (32), as well as the animal’s speed, which is known to substantially influence global hippocampal activity (61, 62), can increase global synchrony and introduce spurious correlations. It is only by factoring out all these known sources of covariability, compactly captured by spike synchrony (35), that the fine structure of pairwise cell interactions can be revealed. To reliably detect such interactions, we developed a novel statistical test rooted in the maximum entropy framework (34).

When applying our detection method to tetrode recordings of hundreds of isolated units in dorsal hippocampus of freely behaving rats (37, 38), we found stark differences between familiar and novel environments, especially in the EE subnetwork. In particular, we found increased interactions among putative pyramidal neurons in novel environments. Furthermore, we detected increased interactions between excitatory and inhibitory cells in novel environments. This effect was not explained by higher reliability of direct excitatory-inhibitory connections (47). It has long been known that inhibition is generally weaker in a novel vs. a familiar environment (15– 17), which has been interpreted as a potential mechanism for enhancing learning by promoting synaptic plasticity in excitatory neurons (15, 43). Nonetheless, given that the null models capture both single cell average activity and population synchrony for each environment separately, it is unlikely that this observation can directly account for our results. Instead, our observations in the novel environment are likely to derive from an increased excitability at the dendritic level of pyramidal cells, an effect that has been observed experimentally (63) and has theoretically been shown as necessary for place field formation and stabilization (19).

Our key statistical observation could be distilled into one simple principle: a monotonic nonlinear dependence of the interaction probability on place field overlap for positive interactions among excitatory cells. This effect was observed across experience, but was more prominent during novelty. We analysed the neural coding implications of the inferred interaction structure using stimulus-dependent pairwise maximum entropy models (53). We found that data-like interactions offered improvements in spatial information content and decoding. Coding advantages were higher during novelty: this observation argues for a mechanism employed by CA1 networks to cope with worse quality input from CA3 (13) and MEC (20, 21) during novelty. We also found that data-like interactions improved stimulus discriminability, corroborating previous findings (30). Moreover, our results explain why disrupting correlations between hippocampal neurons leads to decreased decoding accuracy (10).

Efficient coding in the place cell network yields optimal solutions in which similarly tuned neurons have a higher probability of interacting positively. This is especially prominent for lower-quality inputs in the novel environment, where the predicted relation between interaction probability and tuning similarity is clearly nonlinear, as observed in data. Simulated networks where this observed relationship is elevated to an actual connectivity rule show that, *(i)*, the observed relationship is sufficient to improve population spatial coding, and *(ii)*, the resulting network topology shows clear small-world fingerprints (52, 64). While our results point towards small-worldness as one consequence of the particular connectivity rule that may be employed in the hippocampus (65), they do not provide any evidence that small-world networks have intrinsic coding benefits *per se* (66, 67). Further work is needed to clarify the relationship between coding and small-worldness and to experimentally probe whether small-world architecture is common in networks that need to process noisy inputs.

Even though inferred pairwise interactions do not necessarily reflect underlying synaptic connectivity directly (68), together with the neuron tuning function they offer an accurate statistical description of a neural population output (11, 69, 70). Moreover, pairwise interactions can be studied using well established tools from information theory, which critically rely on the differentiation between stimulus selectivity overlap and network interactions to assess the amount of information that a population carries about a stimulus (29). We derived and tested the efficient coding hypothesis for a network of interacting place cells, by maximizing the mutual information between the animal’s location (the stimulus) and the population response, while holding individual cell tuning and overall firing rate fixed. We found that network interactions adapt to different levels of input quality by employing different interaction vs. tuning similarity strategies. In particular, for low input quality (i.e., at low signal-to-noise ratio mimicking the novel environment) optimal network interactions are strongly aligned with the tuning similarity of the interacting cells. When input quality is higher (i.e., at higher signal-to-noise ratio mimicking the familiar environment), this relation weakens yet remains detectable. These optimality predictions closely resemble the data, suggesting that the CA1 circuit is close to an optimal operating regime across experience. As far as we know, this study is the first empirical test of the efficient coding hypothesis applied to network interactions, as proposed by previous theoretical work (29).

Theory predicts the inversion of the relative contribution of optimal interaction and tuning at very high signal-to-noise ratios (29). This causes the neural population to decorrelate its inputs, a regime that is characteristic for coding in the sensory periphery. While our numerical simulations reproduce this decorrelation regime of efficient coding at very high signal-to-noise ratio inputs, our inferences and data analyses suggest that it is not relevant for the hippocampal place code. This is likely because the overall noise levels are higher in the spatial navigation circuits compared to the sensory periphery, and partially because of the intrinsic differences in the statistics of the signal to be encoded (position vs. natural images). Further work is needed to quantitatively relate the experimentally measured noise in CA1 inputs and responses to the effective “input quality” parameter that enters our predictions.

Are there previous reports where efficient coding predictions do not lead to decorrelation? A classic analysis in the retina correctly predicted that the receptive fields should lose their surrounds and switch to spatial averaging at low light (71). A detailed study of retinal mosaics suggested that even during day vision receptive field centers of ganglion cells should (and do) overlap, increasingly so as the noise increases, leading to a residual redundancy in the population code (72, 73), as reported (74). These findings support a more nuanced view of retinal coding (75) than the initial redundancy reduction hypothesis (76), precisely because they take into account the consequences of noise in the input and circuit processing (77– 79). A recent study in fly vision focused on an interaction between two identified neurons, to find that its magnitude increased as the visual input became more and more noisy, as theoretically predicted by information maximization (80). Psychophysics of texture sensitivity that arises downstream of the primary visual cortex further suggested that the relevant neural mechanisms operate according to the efficient coding hypothesis, yet in the input-noise-dominated regime where decorrelation is not optimal (81). In light of these examples and our results, efficient coding—understood more broadly as information maximization (82) rather than solely in its noiseless decorrelating limit—should be revisited as a viable candidate theory for representations in the central brain. More generally, our approach enables a synergistic interplay between statistical analysis, information theory, graph theory and traditional neural coding, and opens new ways for investigating neural coding during complex/naturalistic behavior in other systems.

## Materials and Methods

### A. Experimental procedures

#### Datasets and Subjects

We analyzed data from two previously published datasets (37, 38). All procedures involving experimental animals were carried out in accordance with Austrian animal law (Austrian federal law for experiments with live animals) under a project license approved by the Austrian Federal Science Ministry. Four adult male Long-Evans rats (Janvier, St-Isle, France) were used for the experiments in (38). We used two wildtype littermate control animals, generated by breeding two DISC1 heterozygous Sprague Dawley rats from (37). Rats were housed individually in standard rodent cages(56×40×26 cm) in a temperature and humidity controlled animal room. All rats were maintained on a 12 hr light/dark cycle and all testing performed during the light phase. Food and water were available *ad libitum* prior to the recording procedures and bodyweight at the time of surgery was 300-375 g.

#### Surgery

The first 4 animals (38) were implanted with microdrives housing 32 (2×16) independently movable tetrodes targeting the dorsal CA1 region of the hippocampus bilaterally. Each tetrode was fabricated out of four 10 um tungsten wires (H-Formvar insulation with Butyral bond coat California Fine Wire Company, Grover Beach, CA) that were twisted and then heated to bind them into a single bundle. The tips of the tetrodes were then gold-plated to reduce the impedance to 200-400 kU. During surgery, the animal was under deep anesthesia using isoflurane (0.5%–3% MAC), oxygen (1-2l/min), and an initial injection of buprenorphine (0.1mg/kg). Two rectangular craniotomies were drilled at relative to bregma (centered at AP =-3.2; ML = *±*1.6), the dura mater removed and the electrode bundles implanted into the superficial layers of the neocortex, after which both the exposed cortex and the electrode shanks were sealed with paraffin wax. Five to six anchoring screws were fixed on to the skull and two ground screws (M1.4) were positioned above the cerebellum. After removal of the dura, the tetrodes were initially implanted at a depth of 1-1.5 mm relative to the brain surface. Finally, the micro-drive was anchored to the skull and screws with dental cement (Refobacin Bone Cement R, Biomet, IN, USA). Two hours before the end of the surgery the animal was given the analgesic Metacam (5mg/kg). After a one-week recovery period, tetrodes were gradually moved into the dorsal CA1 cell layer (stratum pyramidale).

The last two animals (37) were implanted with microdrives housing 16 independently movable tetrodes targeting the right dorsal CA1 region of the hippocampus. Each tetrode was fabricated out of four 12 um tungsten wires (California Fine Wire Company, Grover Beach, CA) that were twisted and then heated to bind into a single bundle. The tips of the tetrodes were gold-plated to reduce the impedance to 300-450 kΩ. During surgery, the animal was under deep anesthesia using isoflurane (0.5-3%), oxygen (1-2 L/min), and an initial injection of buprenorphine (0.1 mg/kg). A rectangular craniotomy was drilled at -3.4 to -5 mm AP and -1.6 to -3.6 mm ML relative to bregma. Five to six anchoring screws were fixed onto the skull and two ground screws were positioned above the cerebellum. After removal of the dura, the tetrodes were initially implanted at a depth of 1-1.5 mm relative to the brain surface. Finally, the microdrive was anchored to the skull and screws with dental cement. Two hours before the end of surgery the analgesic Metacam (5 mg/kg) was given. After a one-week recovery period, tetrodes were gradually moved into the dorsal CA1 cell layer.

After completion of the experiments, the rats were deeply anesthetized and perfused through the heart with 0.9% saline solution followed by a 4% buffered formalin phosphate solution for the histological verification of the electrode tracks.

#### Behavioral procedures

Each animal was handled and familiarized with the recording room and with the general procedures of data acquisition. For the first 4 animals (38), four to five days before the start of recording, animals were familiarized at least 30 min with a circular open-field environment (diameter = 120 cm). On the recording day, the animal underwent a behavioral protocol in the following order: exploration of the familiar circular open-field environment (40 mins), sleep/rest in rest box (diameter =26cm, 50 mins). Directly after this rest session the animals also explored a novel environment for an additional 40 min and rested after for 50 mins. The novel environment recordings were performed in the same recording room but in an enclosure of a different geometric shape but similar size (e.g., a square environment of 100cm width). The wall of both the familiar and novel environment enclosures was 30cm in height, which limited the ability of the animal to access distal room cues. In addition, in two animals a 50 mins sleep/rest session was performed before the familiar exploration.

For the last 2 animals (37), two to three days before the start of recording, animals were familiarized with a circular open-field environment (diameter = 80 cm). On the recording day, the animal underwent a behavioral protocol in the following order: 10 min resting in a bin located next to the open-field environment, exploration of the familiar open-field environment (20 min), sleep/rest in the familiar open-field environment (20 min), exploration of a novel open-field environment (20 min), sleep/rest in the novel open-field environment (20 min). Whilst the familiar environment was kept constant, the novel environment differed on every recording day. The novel open-field arenas differed in their floor and wall linings, and shapes. The recordings for the familiar and novel conditions were performed in the same recording room.

During open-field exploration sessions, food pellets (MLab rodent tablet 12mg, TestDiet) were scattered on the floor to encourage foraging and therefore good coverage of the environment.

#### Data Acquisition

A headstage with 64 or 128 channels (4 × 32 or 2 × 32 channels, Axona Ltd, St. Albans, UK) was used to preamplify the extracellular electric signals from the tetrodes.

Wide-band (0.4 Hz–5 kHz) recordings were taken and the amplified local field potential and multiple-unit activity were continuously digitized at 24 kHz using a 128-channel (resp. 64-channels) data acquisition system (Axona Ltd St. Albans, UK). A small array of three light-emitting diode clusters mounted on the preamplifier headstage was used to track the location of the animal via an over-head video camera. The animal’s location was constantly monitored throughout the daily experiment. The data were analyzed offline.

### B. Data Processing

#### Spike sorting

The spike detection and sorting procedures were performed as previously described (83). Action potentials were extracted by first computing power in the 800-9000 Hz range within a sliding window (12.8 ms). Action potentials with a power >5 SD from the baseline mean were selected and spike features were then extracted by using principal components analyses. The detected action potentials were segregated into putative multiple single units by using automatic clustering software (http://klustakwik.source-forge.net/). These clusters were manually refined by a graphical cluster cutting program. Only units with clear refractory periods in their autocorrelation and well-defined cluster boundaries were used for further analysis. We further confirmed the quality of cluster separation by calculating the Mahalanobis distance between each pair of clusters (39). Afterwards, we also applied several other clustering quality measures and selected only cells which passed stringent measures. In particular we implemented: isolation distance and l-ratio (40), ISI violations (41) and contamination rate. We employed the code available on Github: https://github.com/cortex-lab/sortingQuality. The criteria for the cells to be considered for analysis were the following:

- Isolation distance > 10−th percentile
- ISI violations *<* 0.5
- contamination rate *<* 90−th percentile

Periods of waking spatial exploration, immobility, and sleep were clustered together and the stability of the isolated clusters was examined by visual inspection of the extracted features of the clusters over time. Putative pyramidal cells and putative interneurons in the CA1 region were discriminated by their autocorrelations, firing rate, and waveforms, as previously described (Csicsvari et al., 1999a).

#### Data inclusion criteria

We set a minimum firing rate of > 0.25 Hz on average, across both familiar and novel environments. The final dataset consisted of 294 putative excitatory and 128 putative inhibitory cells across 6 animals. Considering only pairs of units recorded on different tetrodes, the dataset includes a total of 9511 excitatory-excitatory (EE) pairs, 7848 excitatory-inhibitory (EI) and 1612 inhibitory-inhibitory (II) pairs.

Spiking data was binned in 25.6 ms time windows, reflecting the sampling rate for positional information. We excluded bins where:

- the animal was static (speed *<* 3cm/s)
- sharp-wave ripple oscillatory activity was high, i.e. periods with power in the band 150 ∼ 250 Hz in the top 5th percentile (83, 84)
- theta oscillatory activity was particularly low, with power in the band 5 ∼ 15 Hz in the lowest 5th percentile; it is known that hippocampal theta oscillations support encoding of an animal’s position during spatial navigation and reduces overall synchrony of population (85, 86).

#### Theta phase detection and data binning in theta cycles

MN: we are not talking about this in the paper. Exclude?

### C. Null model of population responses and detection of excess correlations

#### Maximum entropy null model

We construct a null model for population responses that takes into account the position of the animal, **s** and the population synchrony, *k* = ∑_*i*_ *x*_*i*_, but is otherwise maximally variable. We use this model to generate a large ensemble of surrogate datasets, that match the data with respect to tuning but without additional noise correlations. Using these surrogates allow us to estimate an empirical distribution of (total) pairwise correlations under the null model, which we then compare to data.

Under the assumption that spike counts have mean *λ*(**s**, *k*) with Poisson noise, the distribution of the joint neural responses under the null model factorizes as:

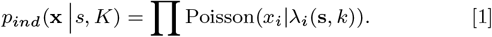

One important caveat is that the population synchrony depends on the neural responses themselves, which introduces the additional constraint that *k* = ∑_*i*_ *x*_*i*_ for each of these surrogate draws, something that we enforce by rejection sampling (87). The only remaining step is to estimate the tuning function of each cell, *λ*_*i*_(**s**, *k*), which we achieve using a nonparametric approach based on Gaussian Process (88) priors.

#### Tuning function estimation

Here we briefly describe the key steps of the approach, and refer the reader to (89) for further details. The data is given as *T* input pairs, *𝒟* = {***x***_*i*_, *y*_*i*_}_*i*=1,2,…,*T*_, where **x**_*i*_ denotes the input variables, defined on a 3−dimensional lattice for the 2*d−*position of the animal in the environment and population synchrony, defined as 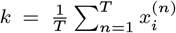; and *y*_*i*_ denotes spike counts in the corresponding time bin (*dt* = 25.6ms).

Neural activity is modeled as an inhomogeneous Poisson process with firing rate dependent on input variables, *λ*(**x**_*i*_). We use a Gaussian Process (GP) prior to specify the assumption that the neuron’s tuning is a smooth function of the inputs, with an exponential link function, *f* = log *λ, f* ∼ 𝒢 𝒫 (*µ, k*), with mean function *µ*(·) and covariance function *k*(*·,·*). In particular, we use a product of squared exponential (SE) kernels for the covariance function:

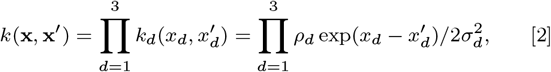

This allows the prior covariance matrix to be decomposed as a Kronecker product *K* = *K*_1_ ⊗ *K*_2_ ⊗ *K*_3_, dramatically increasing the efficiency of the fitting procedure (90).

The parameters *θ* = {*µ, ρ, σ*} are fitted from data by maximizing the marginal likelihood of the data given parameters. Given estimated parameters, 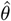, we infer the predictive distribution 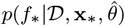 for a set of input values **x**_∗_ (defined below). This distribution can be computed by marginalizing over **f** :

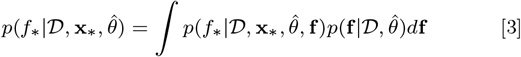

This distribution is intractable, but can be approximated by using a Laplace approximation for 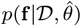 so that ultimately 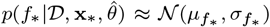. Finally, thanks to the exponential link function, the inferred firing rate of an individual input point *λ*(**x**_∗_) = exp(*f*_∗_) is log-normally distributed, whose mean and variance can be easily computed as:

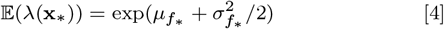

and

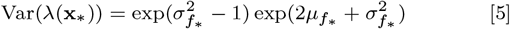

We chose input points **x**_∗_ = (**s**, *k*) that corresponded to the binned 2D location **s** of the animal (5cm bins) and binned population synchrony *k* (10 equally weighted bins, each containing 10% of the data, i.e. the bin edges correspond to the (0th, 10th …, 100th) percentiles).

#### Generating surrogate data

At each moment in time, given the position **s** and population synchrony *k*, the GP tuning estimate provides a distribution over possible firing rates for cell *i, λ*_*i*_(**s**, *k*), as a log normal distribution, 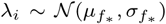. This captures uncertainty about the tuning of the cell, given the data. We generate surrogate spike counts in two steps. First, we sample the mean firing from this *p*(*λ*_*i*_ |*s, K*) distribution. Second, for each *λ*_*i*_ sample, we draw the corresponding spike count from Poisson(*λ*_*i*_). Applying this procedure for all cells and all time points generates a surrogate dataset from the unconstrained null model. We enforce the con straint ∑_*i*_*x*_*i*_ = *k* by discarding and redrawing samples that do not satisfy it. In rare cases (less than 2% of data), it was not possible to replicate the desired *k* statistic, i.e. achieving the desires *k* required more than 500 re-samplings. Such time bins were excluded from subsequent analysis (both for for real data and all surrogates). We generate a total of 1000 surrogate datasets.

#### Inference of excess correlations

We use the pairwise correlations between neural responses as the test statistic and compare it to the distribution of pairwise correlations expected under the null model that assumes that the firing rate of cells is only driven by the stimulus and the synchrony of the population, without further pairwise interactions.

Given the Pearson correlation coefficient between the activities of cells *i* and *j* computed on real data, *c*_*ij*_, and 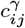 the same quantity computed on a surrogate dataset 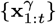 for *γ* = 1, 2, … 1000. We define the quantity we refer to as “excess correlations” as:

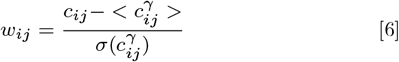

where *< · >* denotes the sample average and *σ* the sample standard deviation of 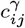. Assuming that the 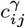 distribution is normal, this quantity is closely related to confidence bounds, and p-values (via the error function). An excess correlation is deemed significant if |*w*_*ij*_|> 4.5, which corresponds to a p-value threshold of *p* = 0.05 with a Bonferroni correction for the 7500 multiple comparisons.

#### Validation

To validate our method, we construct an artificial dataset with know interactions, by sampling from a coupled stimulus de pendent MaxEnt model. We consider *N* = 50 neurons and binary activations **x** = (*x*_1_, … *x*_*N*_) ^⊤^ for any given time window. The distribution of responses **x** given a location-stimulus *s* and synchrony level *k* is

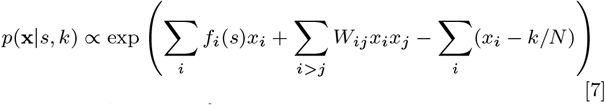

where *s* ∈ {*s*_1_, …, *s*_*K*_} is a spatial position chosen from a set of discrete locations uniformly spaced in the environment, and the feedforward input to each cell, *f*_*i*_ = *f*_*i*_(*s*), is as described in methods subsection (D). We try to match the general statistics of the data as closely as possible. In particular, we match the true time-dependent occupancy, *s*_*t*_, observed in a 20 minutes exploration session, and the corresponding time-dependent synchrony observed in the same session, *k*_*t*_, by sampling one population activity vector (after adequate burn-in time) at each time point **x**(*t*) ∼*P* (**x** |*s*_*t*_, *k*_*t*_) using Gibbs sampling (91).

Given this artificial dataset, we analyze it with the same processing pipeline that we use for the neural recordings and compare the estimated interactions *w*_*ij*_ with the ground truth couplings *W*_*ij*_, which are randomly and independently drawn from 𝒩 (0, 1). Furthermore, we generate data with the same constraints but without any interactions. We asses the ability of our statistical test to detect true interactions using the receiver operating characteristic (ROC), and estimate false positive rates for our statistical test.

### D. Hippocampal population responses with adjustable network structure

#### Stimulus dependent MaxEnt model

In order to explore the effects of the noise correlation structure on the coding properties of the hippocampal system, we employed a statistical model of the collective behavior of a population of place cells that allowed us to vary the couplings among cells while keeping fixed the output firing rate. A similar, stimulus dependent maxent model was introduced in (53), and more recently was used in (11) to prove that correlation patterns in CA1 hippocampus are not due to place encoding only, but also to internal structure and pairwise interactions. Our model includes spatially-selective inputs with adjustable strength, *h*, and noise correlations modelled as a matrix **W** describing the strength of interaction between cell pairs. Additionally, we constrained average population firing rates to be the same for each possible choice of *h* and **W**, as a way of implementing metabolic resource constraints.

More specifically, consider *N* neurons with binary activations **x** = (*x*_1_, … *x*_*N*_) ^⊤^. The distribution of responses **x** given a location-stimulus *s* we considered is

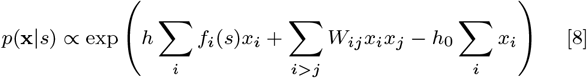

where *s* ∈ *{s*_1_, …, *s*_*K*_} is a spatial position chosen from a set of discrete locations uniformly spaced in the environment (the unit square, [0, 1] *×* [0, 1]). The feedforward input to each cell, *f*_*i*_ = *f*_*i*_(*s*), is modelled as a 2 − D Gaussian bump with continuous boundary conditions, mean randomly drawn from a uniform on [0, 1] *×* [0, 1] and fixed covariance 0.1𝕀. The parameter *h*_0_ allows us to fix the average population firing rate to 20% of the population size, and is found by grid optimization. Once the input tuning *f*_*i*_ is fixed for each cell, we select the connections *W*_*ij*_ for each cell pair by sampling from the data-inferred excess correlations of cell pairs with similar tuning similarity, and then scaling according to the results found during method validation (Fig S 1). We did so separately for familiar and for novel environments. Finally, we fix the appropriate parameter *h*, separately for familiar-like and novel-like connections, by matching single neurons marginal statistics. We utilized three measures: single cell spatial information, sparsity and gain, which are described in detail in Methods subsection (E).

#### Optimization of connections for fixed input and fixed firing rate

Given *h*, {*f*_*i*_(·)}, we optimize the connections **W** so as to maximize the mutual information between population activity and spatial position, 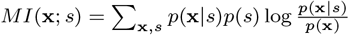, via Sequential Least SQuares Programming (SLSQP) (92). We further constrain the population average firing to 20% of the nural population, and each *W*_*ij*_ is restricted to lay in [−1, 1]. Both reflect biological resource constraints on the optimal solution.

Most simulations use *N* = 10 neurons, which allows the mutual information to be computed in closed form (by enumerating all possible patterns). Reported estimates are obtained by averaging across 1000 randomly initialized networks (different *f*_*i*_(·) centers, and initial conditions for the optimization). To ensure that our results generalize to large networks, we also performed limited numerical simulations for *N* = 20 (only for *h* = 2 and *h* = 4, averaging over 10 networks.

#### Optimal coding for large networks

The exact computation of the mutual information *MI*(**x**; *s*) is very resource intensive and only applicable to small networks (*N* ≤ 20). To investigate the effects of noise correlations at larger scales we need to rely on efficient approximations. The mutual information between population binary responses **x** and location-stimulus *s* can be written as

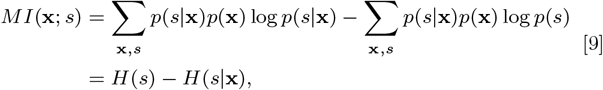

where *H* denotes (conditional) entropy. Assuming that *p*(*s*) is a uniform distribution over stimuli, we have *H*(*s*) = 2 log *B*, where *B* is the number of bins used to discretize each dimension of the 2 dim environment. We generally use *B* = 16. The challenge is to compute *H*(*s* |**x**). For a given **x**, denote with *ĥ*(**x**) := −∑_*s*_*p*(*s* |**x**) log *p*(*s* |**x**). Then we have:

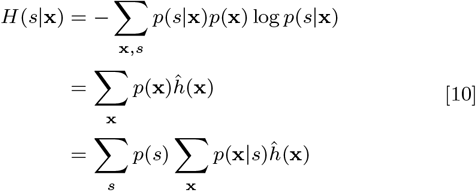

We used the last expression and estimated *H*(*s*|**x**) by drawing 10^6^ samples from *p*(**x**|*s*) for each stimulus *s* using Gibbs sampling (91).

We reported the estimated average across stimuli and confidence intervals in the figures. The quantity *ĥ*(**x**) := −∑_*s*_*p*(*s* |**x**) log *p*(*s* |**x**) is the entropy of the posterior distribution on stimuli given a certain binary vector. The main obstacle to computing *ĥ* is that, for each stimulus *s*, we need to know the proportionality constant *Z*_*s*_ = ∑_**x**_ *p*(**x** |*s*) (i.e. the partition function), that makes the probability (8) sum up to 1. We computed *Z*_*s*_ exhaustively for *N* ≤ 20 by enumerating all the possible binary vectors. For *N* ≥ 20 we estimated it using a simple Monte Carlo method by randomly drawing 10^9^ independent *N* −dim binary samples for each stimulus, and then regularizing by applying a mild 2D gaussian smoothing (*σ* = 0.5 bins) on the log-transformed *Z*_*s*_ among neighboring stimuli.

#### “Topology” model simulations

We aimed at characterizing the influence of higher order structure on the coding of the network. We used the same model as in eq. [8] with 50 place cells, but allowed connections to be either *−J*, 0 or +*J*, where *J* ∈ [0, 1] is the connection strength. We employed three different strategies to select the units to connect, as described in the main text, based on their tuning similarity. We kept fixed the number of positive (+*J*) and negative (*−J*) couplings to 6% and 3% respectively. For each choice of tuning, connectivity rule and strength *J* we used the parameter *h*_0_ to enforce the population average firing to be 20% of the population size.

### E. Analysis of experimental data

#### Single cell tuning characterization

To describe the tuning properties of single cells we employed several measures:

- gain: peak firing rate over mean, estimated from the tuning function of a cell,
- sparsity: 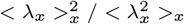, where *λ*_*x*_ denotes the average firing at location *x*, is a measure of how compact the firing field is relative to the recording apparatus (93),
- spatial information: 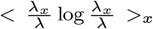, where *λ* =*< λ*_*x*_ >_*x*_, is the leading term of th^*λ*^e MI b^*λ*^etween average spiking and discretized occupancy for small time windows (50, 94).

#### Decoding of spatial position from data

We subdivided the environment in equally spaced 2 −dimensional bins with bin side length of 20 cm. This choice was due to the fact that, to properly estimate the average co-activation of cells one needs many samples and a finer subdivision of the environment made this task extremely difficult. We randomly subdivided the data in two parts, 75% for training and 25% for decoding. With the training data we estimated, for each bin separately, the average activation and the covariance of the neurons activity. With the remaining 25% of the data, we computed for each non-overlapping 10 consecutive 25.6 ms time bins the activation (denoted by population vector or PV) and the covariance (COV). We then simply compared them to all the expected PV and COV measured over the training set in different bins and picked the most similar one in terms of Pearson correlation.

#### PCA, linear separability of pairs of stimuli

We wanted to investigate the linear separability of population responses to different locations. We randomly selected 500 times two distinct locations in the environment and selected all the 250ms population responses in a 10 cm surrounding of the two positions. We then found the best hyperplane that separated the two sets of responses by using a soft-margin linear SVM with hinge loss, and reported the training error. We also computed the principal components of the population responses to both locations together, and reported the variance explained by the first PC.

### F. Network analysis

#### Graph theoretical measures

All the measures were carried out using the library NetworkX (release 2.4) in Python 3.7. We considered unweighted and non directed graphs where each cell was a vertex and an edge connected each cell pair that had a significant interaction (|*w*_*ij*_ |> 4.5). A graph *G* = (*V, E*) formally consists of a set of vertices *V* and a set of edges *E* between them. An edge *e*_*ij*_ connects vertex *v*_*i*_ with vertex *v*_*j*_. The neighbourhood for a vertex *v*_*i*_ is defined as its immediately connected neighbours: *N*_*i*_ = *{v*_*j*_ : *e*_*ij*_ ∈ *E* ∨ *e*_*ji*_ ∈ *E*} and its size will be denoted by *k*_*i*_ = |*N*_*i*_|.

We measured:

1. **Clustering coefficient:** this measure represents the average clustering coefficient of each node, which is defined as the fraction of existing over possible triangles that include that node as a vertex. Formally, the local clustering coefficient *c*_*i*_ for a vertex *v*_*i*_ is given by the proportion of links between the vertices within its neighbourhood divided by the number of links that could possibly exist between them, hence measuring how close its neighbourhood is to forming a clique. If a vertex *v*_*i*_ has *k*_*i*_ neighbours, 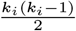 edges could exist among the vertices within the neighbo^2^urhood. Thus, the local clustering coefficient for vertex *v*_*i*_ can be defined as

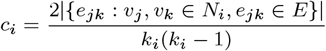

and the average clustering coefficient as

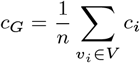
2. **Average shortest path length:** this measure can be computed only if the graph is connected. If not, we computed this measure on the largest connected subgraph.

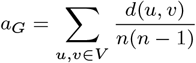

where *u, v* are distinct vertices, *d*(*u, v*) is the shortest path length between *u, v* and *n* is the size of the graph *G*.

#### Triangles analyses

We tested for the over-expression of particular interaction patterns by counting the number of triangles (i.e 3 all-to-all interacting cells) composed by 3 inhibitory cells, 2 inhibitory and 1 excitatory, 1 inhibitory and 2 excitatory or 3 excitatory cells. We tested these counts against the counts from the same networks with shuffled edges. We employed an edge-shuffling procedure that preserved both the total number of edges and the number of incident edges per node, separately for the EE, EI and II subnetworks (i.e. an edge connecting two excitatory cells could be exchanged only with another edge connecting two excitatory edges etc). To do this, we randomly selected two edges of each subnetwork, say *AB* and *CD*. If *A* ≠ *C* ≠ *D* and *B* ≠ *C* ≠ *D* we removed the two edges and inserted the “swapped” ones, *AC* and *BD*. We repeated this procedure 100 times for each subnetwork to yield one shuffled network. We repeated this procedure 1000 times, which gave us a null distribution to test the original counts against. In Supp. Fig. 5 we reported the counts of each pattern, separately for familiar and novel environments, normalized against our null distribution.

#### Betti numbers

We computed the Betti numbers of the clique-complex induced by the graphs. These are distinct from the graphs Betti numbers (95). A clique in a graph is an all-to-all connected set of vertices. The clique complex *X*(*G*) of an undirected graph *G* is an abstract simplicial complex (that is, a family of finite sets closed under the operation of taking subsets), formed by the sets of vertices in the cliques of *G*. Intuitively, the clique-topology can be characterized by counting arrangements of cliques which bound holes. Formally, the dimensions of the homology groups *Hm*(*X*(*G*), ℤ_2_) yield the Betti numbers *b*_*m*_ (95). Given our low connectivity (9%), *b*_*m*_ was almost always zero for *m ≥* 2. On the other side, *b*_0_ simply counts the number of connected components, so in our analysis we focused on *b*_1_. This is the number of cycles, or holes, that are bounded by 1-dim cliques. Graphically, these are 4 edges that form a square, or 5 edges that form a pentagon etc. Notice that 3 edges that form a triangle don’t count towards *b*_1_, because they represent a 2-dim clique (i.e. 3 vertices that are all-to-all connected). This is why a higher clustering coefficient (i.e. more triangles) implies a lower *b*_1_.

## ACKNOWLEDGMENTS

We thank Peter Baracskay, Karola Kaefer and Hugo Malagon-Vina for the acquisition of the data. We thank Federico Stella for comments on an earlier version of the manuscript. MN was supported by European Union Horizon 2020 grant 665385, JC was supported by European Research Council consolidator grant 281511, GT was supported by the Austrian Science Fund (FWF) grant P34015, CS was supported by an IST fellow grant, National Institute of Mental Health Award 1R01MH125571-01, by the National Science Foundation under NSF Award No. 1922658 and a Google faculty award.

## Supplementary figures

**Figure S1.**
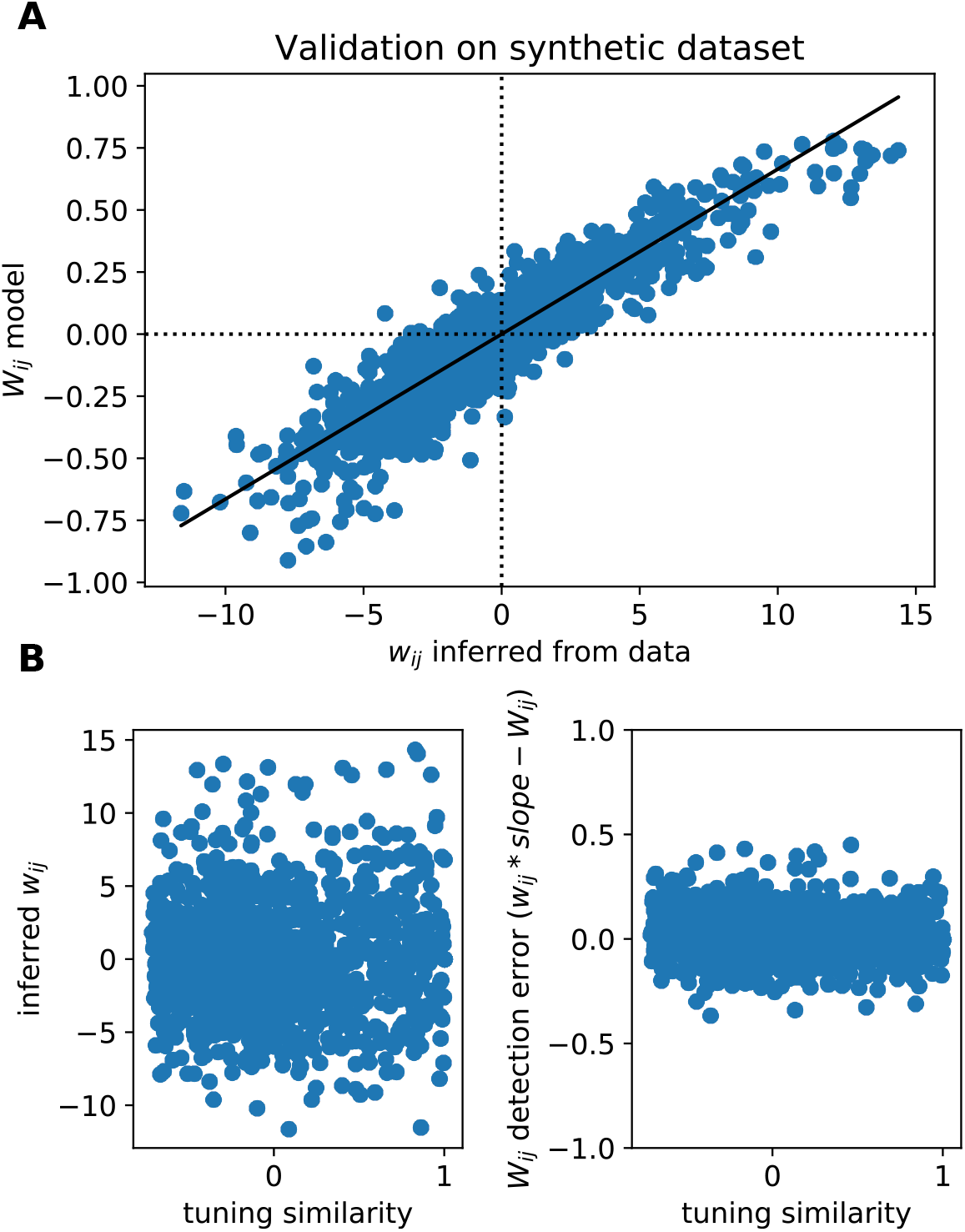
Further data on validation and null model. **(A)** Scatter plot of ground truth *W*_*ij*_ values used in the model for validation vs *w*_*ij*_ inferred from artificial data. **(B)** Left: Scatter of inferred *w*_*ij*_ vs tuning similarity. Notice the absence of bias towards detection for cells with higher or lower tuning similarity. Right: *W*_*ij*_ detection error inferred as the difference between *w*_*ij*_ (scaled by the appropriate slope) and the true *W*_*ij*_. Notice the absence of bias towards highly similarly tuned pairs.

**Figure S2.**
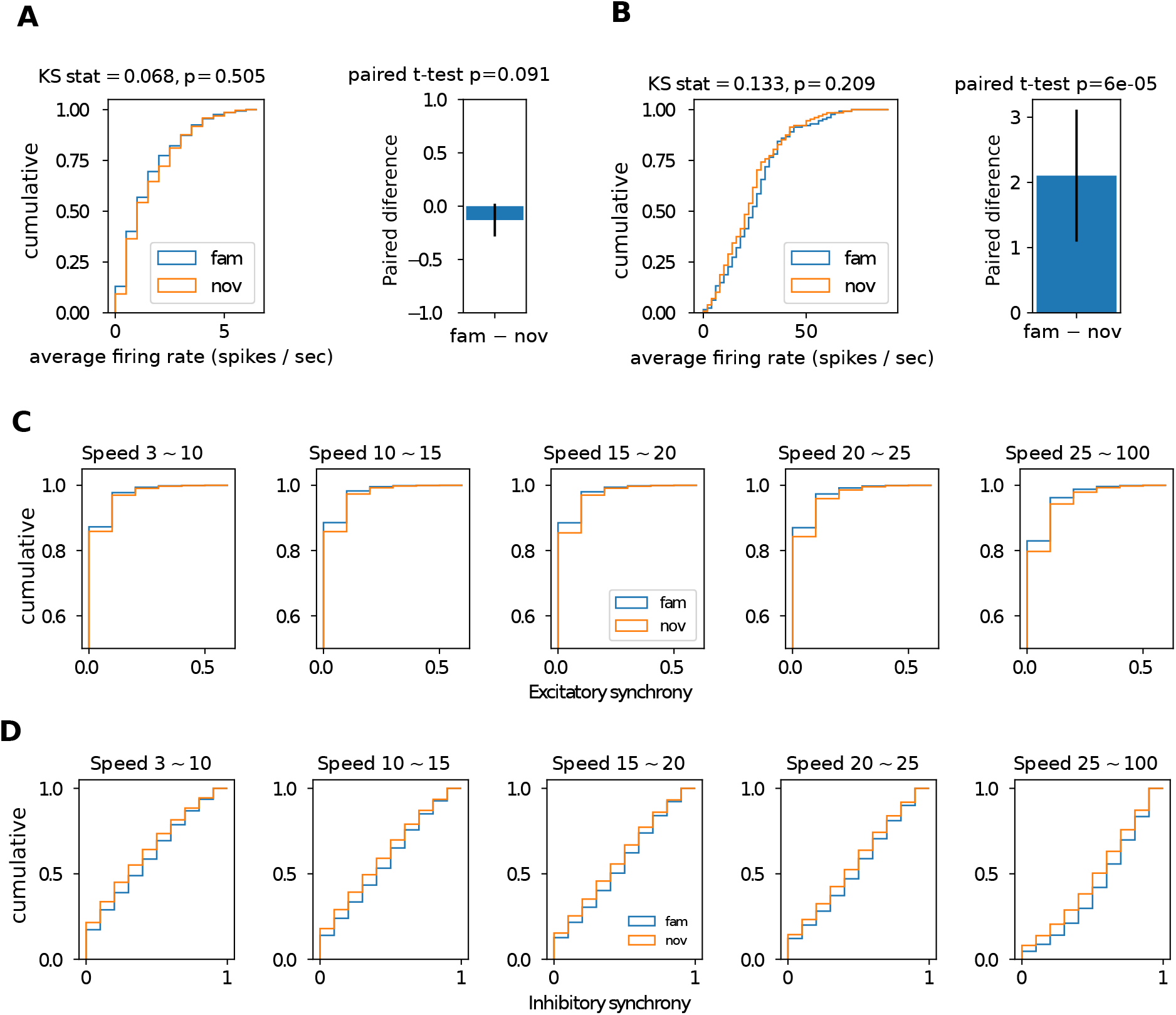
Population marginal statistics. **(A)** Left: distribution of average firing rates of putative CA1 excitatory neurons in familiar (blue) and novel (orange) environment. Right: paired difference across environments (familiar − novel). Error bars represent 95th CI for the mean. **(B)** Same as (A) for putative inhibitory neurons. **(C)** Distribution of synchrony in 25 ms time windows of excitatory neurons for different behavioral speed: [3, 10), [10, 15), [15, 20), [20, 25), [25, 100) cm/sec for familiar (blue) and novel (orange). All KS test *p <* 0.0001. **(D)** Same as (C) for putative inhibitory neurons.

**Figure S3.**
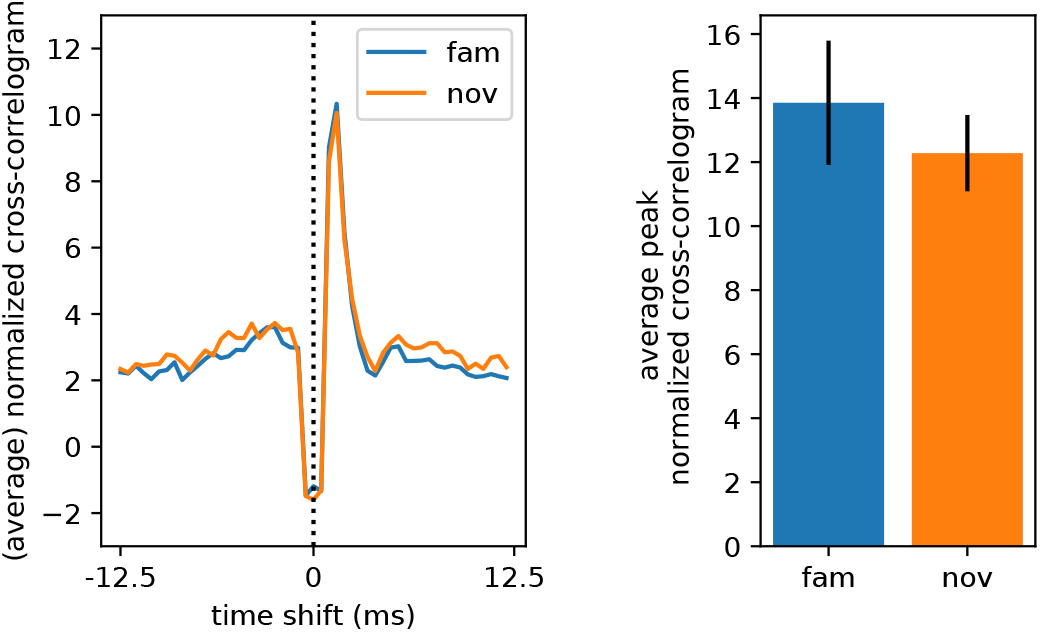
Efficacy of monosynaptic excitatory-inhibitory connections. Left: average normalized cross-correlogram of putative monosynaptically connected excitatory-inhibitory pairs. The cross correlogram was normalized by subtracting the mean and dividng by the STD of cross-correlograms computed on randomly shifted data 100 times. The pairs that had peak (normalized) cross-correlogram > 7*STD* in both environments were labelled as monosynaptically connected (47). Right: average peak of normalized cross-correlogram for familiar and novel environments. Error bars represent 95th CI for the mean. Paired T-test *p* = 0.61.

**Figure S4.**
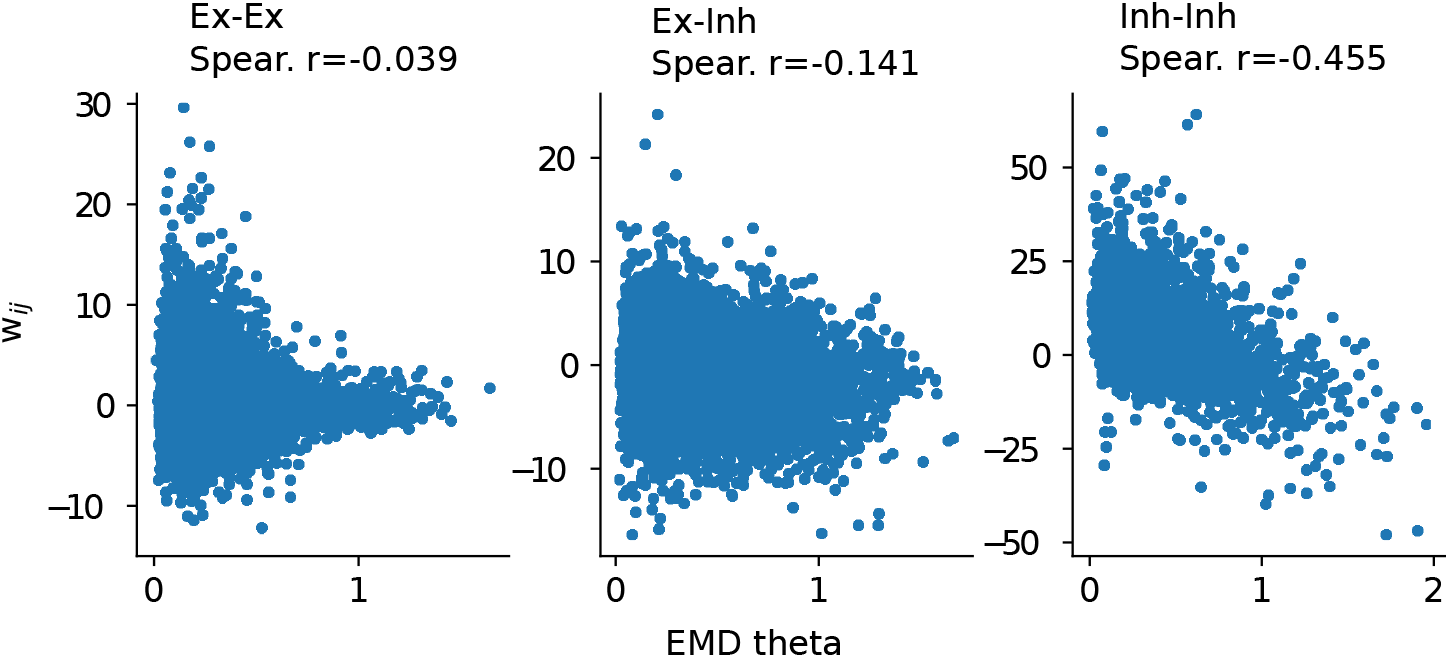
Theta selectivity similarity versus excess correlation. Scatter plot of inferred *w*_*ij*_ vs dissimilarity of theta selectivity measured using an earth mover distance (EMD) among the histograms of preferred theta phases (t-test for Spearman rank correlations: EE *p >* 0.05, EI *p <* 0.001, EE *p <* 0.001).

**Figure S5.**
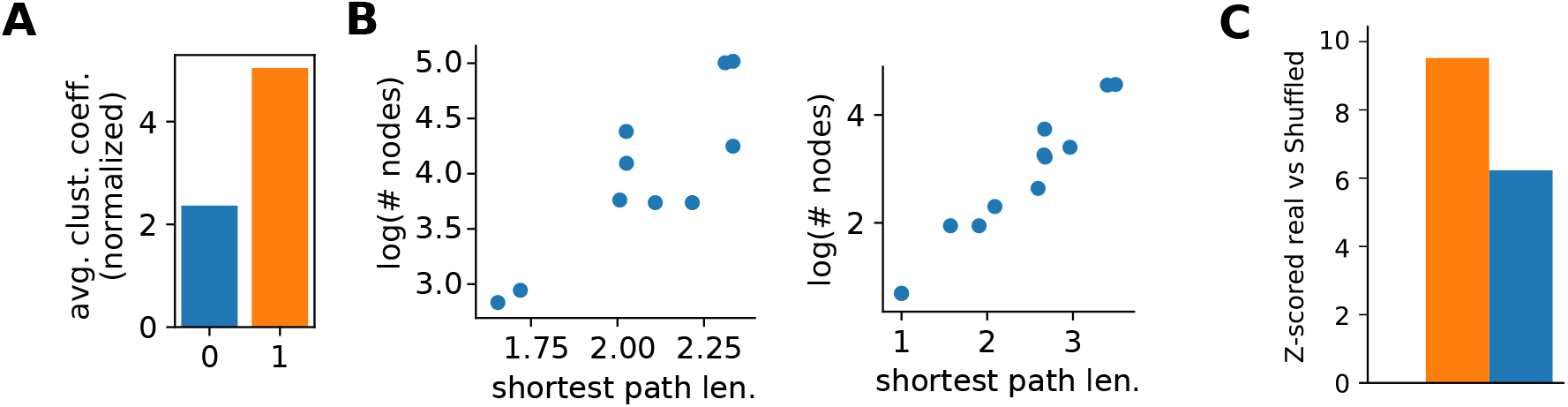
Small worldness of EE subnetwork. **(A)** Average clustering coefficient of excitatory subnetworks normalized against the same values computed on ER random graphs with matching edges density (Fig 2). **(B)** Left: log-nodes number vs shortest path length in the largest connected component of excitatory subnetworks with standard significancy threshold at |*w*| > 4.5 (two dots per animal: familiar and novel). Right: same as left for excitatory subnetworks with higher significancy threshold at |*w*| > 6. **(C)** Overexpression of triangles in real networks against random shuffling of the edges that preserved the number of incident edges onto each single node (see Methods).

**Figure S6.**
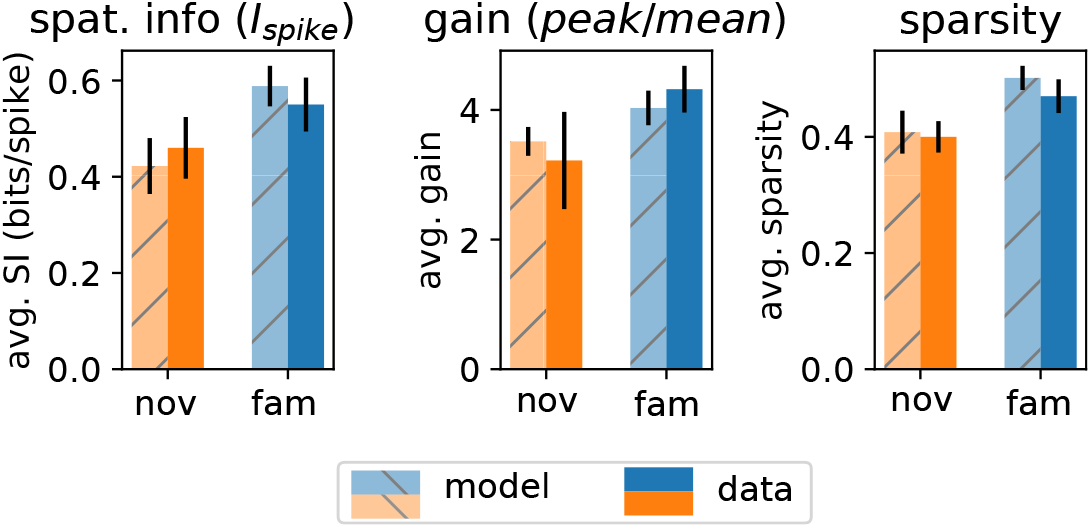
Marginal statistics of place cells in hippocampus match circuit model. The interactions in the model were drawn from the inferred couplings observed in data and rescaled according to Supp. Fig. 1A. Afterwards, we fixed the input strength by picking the parameters that allowed the model to best match the marginal statistics observed in data. All the measures were computed on traditional 2D firing rate maps (see Methods). (left) single cell spatial information, (center) firing rate map gain, measured as peak over mean (right) firing rate sparsity.

**Figure S7.**
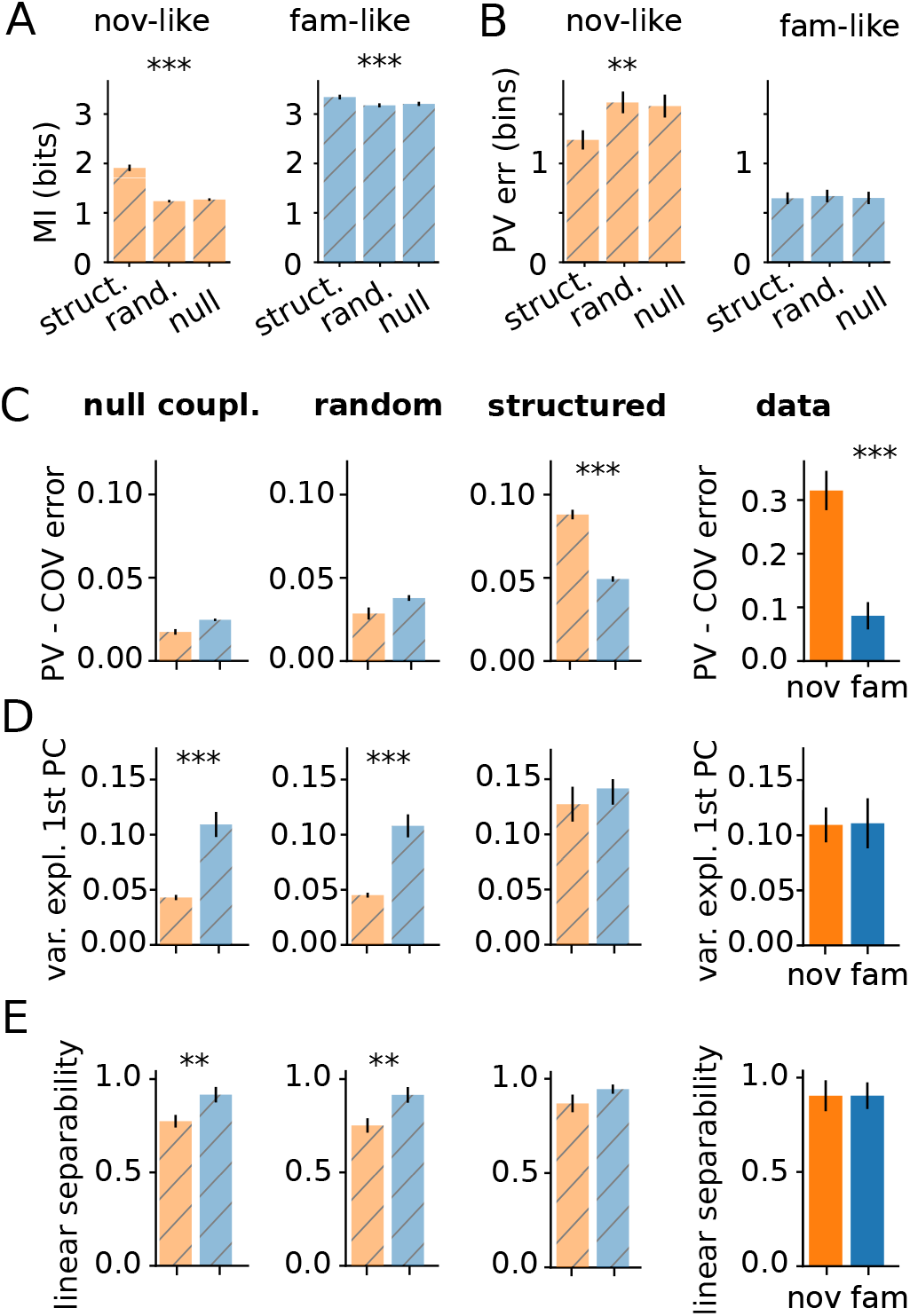
Comparison with null couplings. **(A)** Estimated spatial information (MI; error bar – 99th percentile CI for the mean) using structured, random and null interactions, in the novel-like and familiar-like scenario (see text). Structured interactions significantly increase the spatial information (*p <* 0.001 (***) or *p <* 0.01 (**) under a non-parametric Mann–Whitney U-test). **(B)** Decoding error using a simple population vector approach (PV; error bar – 99th percentile CI for the mean) using structured, random and null interactions, in the novel-like and familiar-like scenario. Structured interactions significantly decrease the average decoding error in novel environments (*p <* 0.01 (**) under a non-parametric Mann–Whitney U-test). **(C)** Improvement in decoding performance by taking into account co-variability of cells (“COV” decoder) relative to a simple population vector (“PV”) decoder, evaluated on 4 ·10^4^ samples). (error bars and significance tests as in B). **(D)** Fraction of variance explained by the first principal component of population vectors for 10^3^ random pairs of locations in the maze. The fraction is unchanged between the novel and familiar environments on structured network and on real data, but differs significantly on the random and null networks (error bars and significance tests as in B). **(E)** Linear separability measured as SVM classification accuracy of random pairs of stimuli (trained on 1000 pairs of same vs. different positions). The separability is unchanged between the novel and familiar environments on structured network and on real data, but differs significantly on the random and null networks.

**Figure S8.**
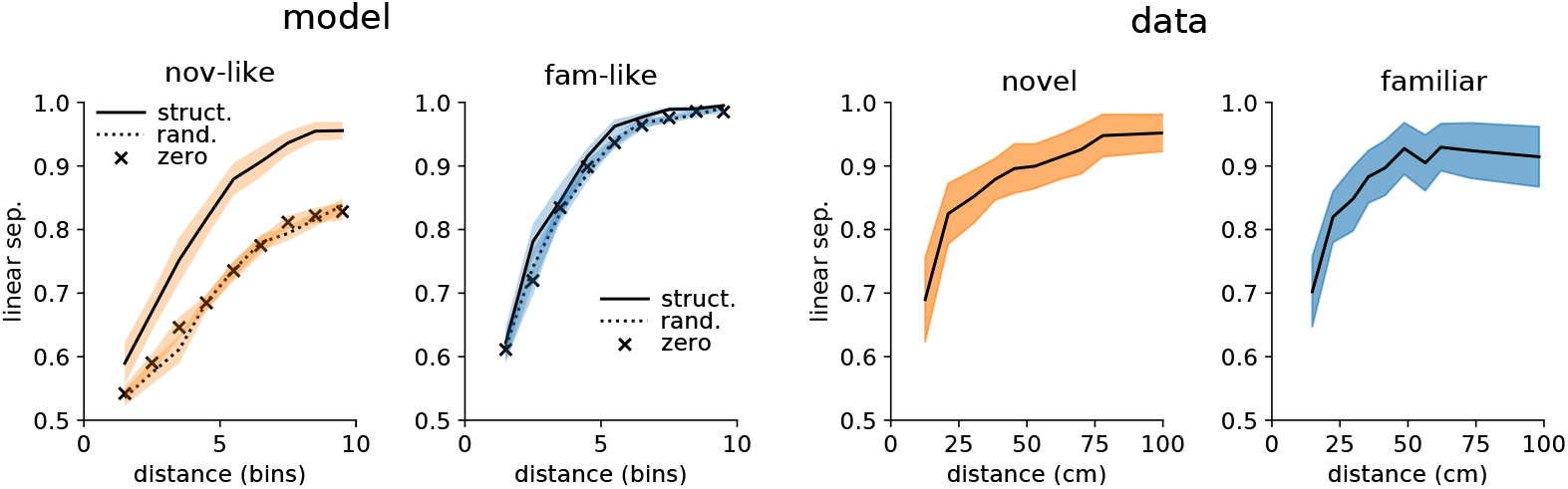
Linear separability as a function of distance. Left: linear separability of responses to stimuli at a given distance for data-like copupling structure (solid line), random connectivity (dotted) or null couplings (x) for novel-like (orange) and familiar-like (blue) input quality. Right: linear separability of responses to stimuli at a given distance for data novel environments (orange) and familiar (blue).

**Figure S9.**
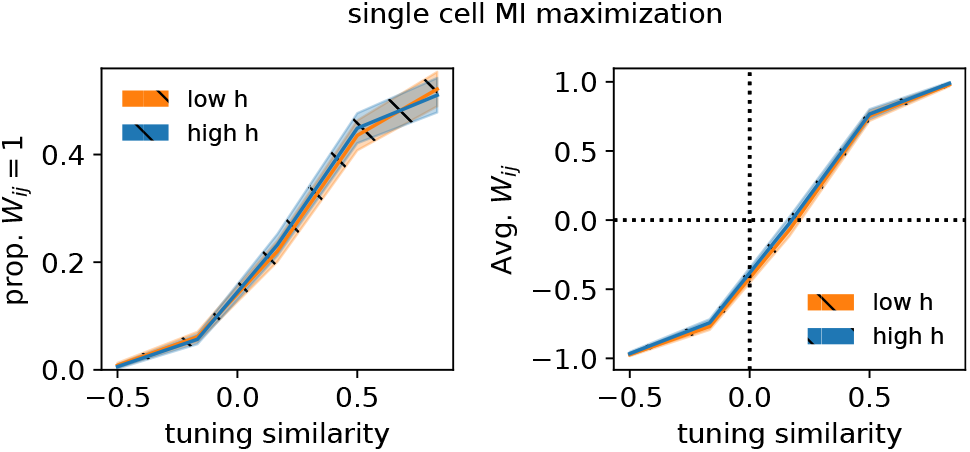
Single cell MI optimization. Optimizing the mutual information between single cells stimulus-dependent (marginalized) activity and location-stimulus led to the same result for each level of input noise – almost linear relation between place field overlap and optimal predicted *W*_*ij*_.

**Figure S10.**
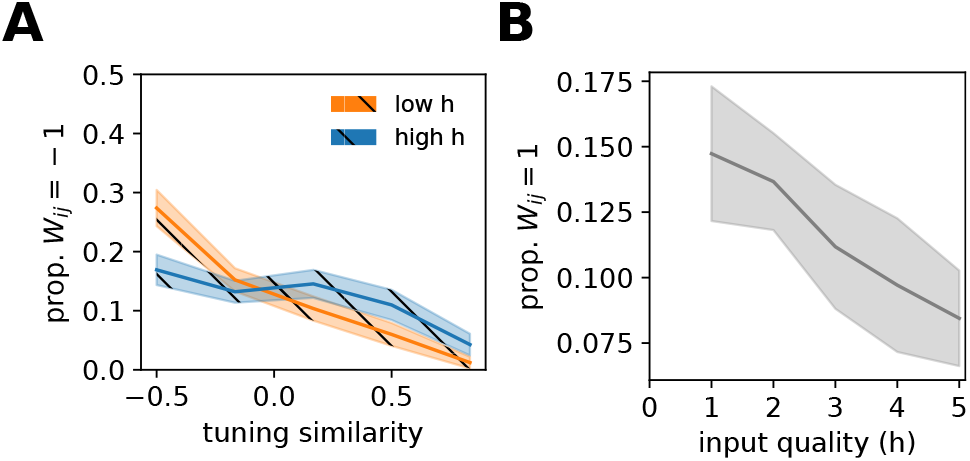
Negatively coupled optimized connections and proportion of strongest. **A** Proportion of cell pairs to reach minimum allowed *W*_*ij*_ as a function of tuning similarity. **B** Proportion of cell pairs that reached maximum *W*_*ij*_ = 1 (after optimization) decreased for increasing input quality *h*.

**Figure S11.**
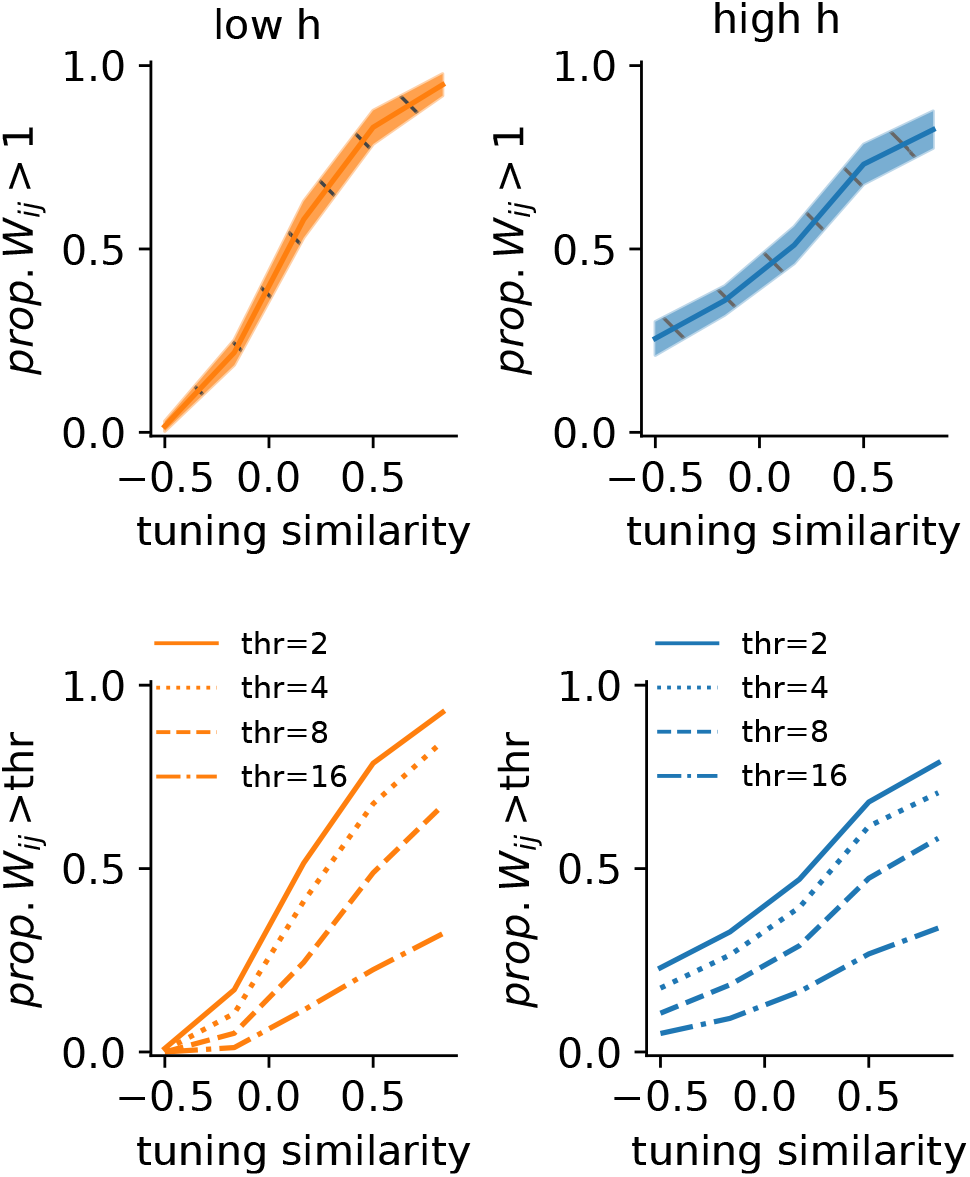
Non constrained maximization does not show nonlinear coupling preferences. Top row: Proportion of couplings that exceed 1 after optimization. Couplings were optimized so to maximize the mutual information between population responses and stimuli. The average population firing rate was constrained but *W*_*ij*_ s were not. Bottom row: mean proportion of couplings that exceed different thresholds also do not show the nonlinear relation we observed in the constrained case.

**Figure S12.**
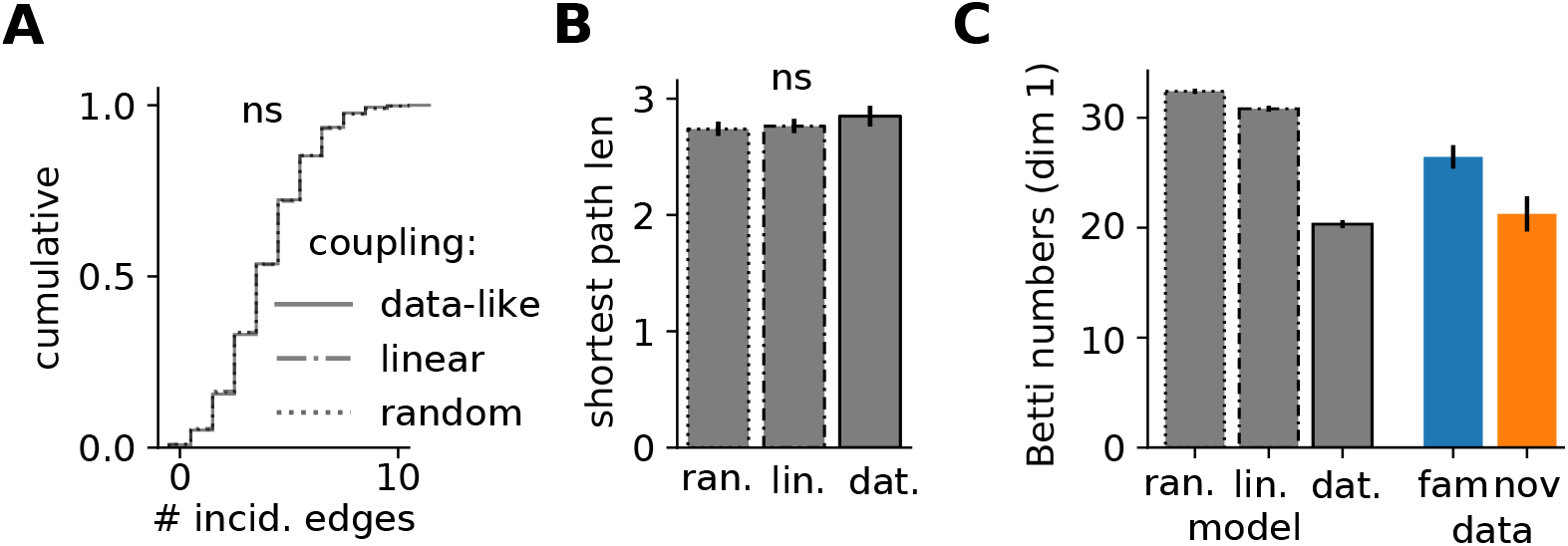
Topology. **(A)** Distribution of incident edges with the three different connectivity-rules. **(B)** Average shortest path length. 1-way ANOVA *p >* 0.05. **(C)** Betti numbers of the clique complex induced by the graph (*b*_1_) for 1-dim holes. Using the data-like nonlinear coupling strategy increased the chance of creating triangles, hence diminishing the number of 1-dim cavities.

**Figure S13.**
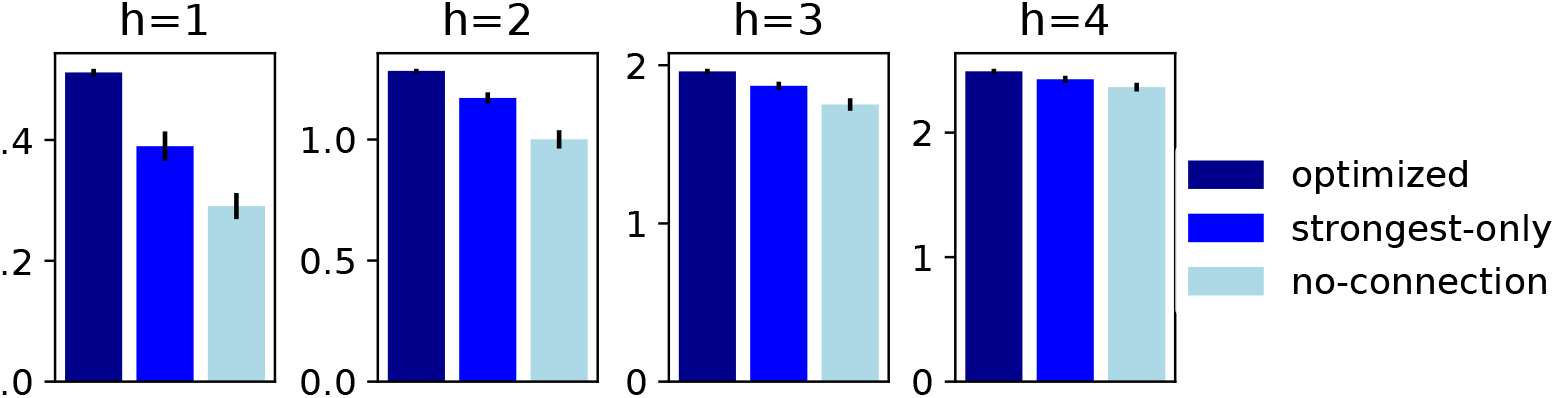
Strongest couplings only. After optimizing the connections *W* (as in Fig. 4), the MI of the fully optimized networks was compared to null couplings and the “strongest only” case, i.e., where every connection |*W*_*ij*_ | *<* 1 was set to 0.

